# Free Energy Landscape of Magnesium Chelation Reveals Dynamic Pre-Chelate Complexes Stabilized by Meta-Sphere RNA-Ion Coordination

**DOI:** 10.1101/2025.05.20.655240

**Authors:** Akhilesh Jaiswar, Raju Sarkar, Avijit Mainan, Rimi Kundu, Susmita Roy

## Abstract

Magnesium ions (Mg²⁺) play a critical role in RNA structure stabilization by forming various coordinated complexes, preferentially interacting with the backbone phosphate groups. Using extensive atomistic and free energy simulations across simple models and RNA structures of varying complexity, we characterized critical components of the RNA-ion-atmosphere. Radial distribution function analysis reveals distinct peak positions for direct (inner) and solvent-separated (outer-sphere) Mg^2+^-phosphate coordination layers, aligning with solution X-ray diffraction data. Addressing forcefield limitations, the free energy calculations quantify the kinetic barriers for Mg²⁺-phosphate binding, benchmarking parameters against ²⁵Mg NMR measurement. Free energy calculations further explore Mg²⁺ chelation with bi-phosphate coordinated Mg^2+^ systems, identifying a dynamic ensemble of pre-chelate complexes, in addition to a chelated and outer-sphere hexa-hydrated state of Mg^2+^. In the pre-chelated states, Mg²⁺ maintains one inner-sphere interaction while simultaneously coordinating with multiple other phosphates in a solvent-separated manner, referred to as meta-sphere coordination. The pre-chelated complexes from different solvents-separated layers undergo a frequent transition and mediate a unique oxygen exchange mechanism between phosphate and water ligands. Insights into the free energy landscape of SAM-I RNA aptamer further emphasize the significance of pre-chelate complexes for complex RNA structure stabilization, where a number of such solvent-separated dynamic phosphate groups are found to influence Mg^2+^-RNA coordination. The comprehensive thermodynamic analysis of Mg²⁺ chelation and quantitative characterizations of various RNA-ion coordination modes, including this new meta-sphere coordination, provides vital insights for advancing RNA modelling and experimental exploration of complex phosphate networks in the RNA structures.

## Introduction

RNA employs various modes of counter-ion interaction to overcome backbone repulsion and achieve its tertiary fold. In a myriad of cellular processes, different ions interact with RNA. Yet, it was the unique potency of Magnesium (Mg²⁺) in stabilizing the tertiary structure of transfer RNA, discovered in the 1970s, that unveiled a far more complex landscape of ion influence on RNA folding.^1–3^ This pivotal discovery underscored the intricate and challenging nature of understanding how ions interact with RNA and their topology. In recent times, the rise of retroviruses, RNA-based vaccines, and the design of RNA biosensors for various medical applications have rekindled interest in the intricate relationship between ions and RNA folding. This renewed focus has sparked a substantial body of literature, exploring the subject through a diverse array of experimental, theoretical, and computational approaches.

Magnesium ions (Mg²⁺), owing to their high charge density, strongly interact with polar solvents such as water. In RNA solutions, the interaction between RNA and Mg²⁺ is modulated by the interactions of Mg²⁺ with water. In bulk solutions, Mg²⁺ predominantly exists in a hexa-hydrated form, surrounded by six water molecules arranged in an octahedral configuration within its primary solvation shell.^4,5^ Pioneering studies by Manning and Record, followed by extensive research and reviews by Draper and colleagues, highlighted the crucial role of this diffuse layer of hexa-hydrated Mg²⁺.^6–10^ These studies attributed its effects to charge screening, which mitigates electrostatic repulsion between RNA backbone segments, a concept widely recognized as Manning’s counter-ion condensation effect.^11^ This diffuse ion-atmospheric layer, formed by various cations, is essential for the stabilization and folding of both secondary and tertiary RNA structures.

As the diffuse layer of Mg²⁺ approaches RNA and forms direct contacts, the hexa-hydrated Mg²⁺ undergoes desolvation, a process that is energetically expensive due to water-exchange kinetics. Crystal structures reveal that such direct contacts typically involve the displacement of 1–3 water molecules from Mg²⁺, forming a partially dehydrated, chelated Mg²⁺ complex.^12–14^ The most common RNA ligands in these interactions are phosphoryl oxygens, purine N7 atoms, keto groups (e.g., O6 of guanosine, O4 of uracil), and ribose 2′-OH groups. Biochemical experiments, including nucleotide analog interference mapping (NAIM), have demonstrated the thermodynamic significance of these direct contacts. Functional substitutions, such as replacing a phosphoryl oxygen-Mg²⁺ interaction with a phosphoryl sulfur-Cd²⁺ interaction, provide further evidence of the role of direct binding in RNA structure and function.^15–17^

Interestingly, in 2002, Anna Marie Pyle and, more recently, Hayes et al. identified a separate class of RNA-Mg²⁺ coordination where Mg²⁺ interacts with RNA without direct contact. Instead, water ligands of the magnesium hexahydrate complex interact with RNA bases and backbone substituents to stabilize specific RNA motifs, a phenomenon referred to as outer-sphere Mg²⁺ interactions.^18,19^ Major groove edges of guanosine, particularly O6 and N7 atoms, are common sites for water-mediated binding. In such cases, Mg²⁺ can often be substituted by exchange-inert mimics like cobalt(III) or osmium(III) hexammine. Studies have demonstrated the role of outer-sphere Mg²⁺ in stabilizing tertiary RNA structures, such as in the BWYV pseudoknot and SARS-CoV-2 RNA pseudoknot, where hydrated Mg²⁺ ions stabilize specific motifs, including tertiary RNA ring structures.^20,21^

Despite the distinction between diffuse, inner-sphere (direct), and outer-sphere Mg²⁺ interactions, the mechanism by which diffuse Mg²⁺ transitions to chelated Mg²⁺ remains poorly understood. This process is crucial for RNA folding and stabilization. Evidence suggests that chelated Mg²⁺ ions, which can simultaneously interact with multiple distant phosphate groups, play a significant role in RNA structure. For instance, in the crystal structure of an intact Group I intron, chelated Mg²⁺ ions were found to stabilize the active site and support ribozyme function.^22^ Similarly, phosphorothioate interference mapping experiments in complex structures like the SAM-I aptamer have revealed buried chelated ions involved in RNA stabilization and ligand binding.^23^

While chelated Mg²⁺ interactions are significant, recent findings emphasize the stabilizing effects of outer-sphere, hydrated Mg²⁺ ions.^19,24^ Despite upgrowing evidence of Mg^2+^-chelation^25,26^, the notion that direct binding is the primary stabilizing force for Mg²⁺ in complex RNA structures remains arguable.^6^ Current understanding suggests that compared to inner-sphere chelated Mg^2+^, the outer-sphere hexa-hydrated Mg^2+^ plays rather a significant role in the stabilizing effect of Mg²⁺ on RNA.^19^ The reason often described as the partial dehydration of a hexa-hydrated Mg²⁺ complex to directly interact with the phosphate groups of RNA is energetically unfavorable, primarily due to the significant entropy loss involved. While this loss of entropy associated with chelation might introduce some electric stress, the strong electrostatic interactions between the chelated ion and multiple phosphate groups could potentially outweigh the entropy loss, making the interaction energetically favorable.^24^ This warrant detailed calculations to characterize the free energy of chelation.

To advance the understanding of ion chelation thermodynamics, appropriate enhanced sampling and free-energy simulation methods are needed. However, the slow exchange rates between Mg²⁺ and water, as well as between Mg²⁺ and phosphate oxygens, present significant challenges for RNA simulations. Villa and coworkers (2012) highlighted limitations in widely used empirical force fields such as AMBER and CHARMM, which underestimate Mg²⁺-water exchange rates by several orders of magnitude.^27^ Although recent updates to RNA force fields have improved RNA structure and dynamics, the critical exchange between inner- and outer-sphere Mg²⁺ ions remains inadequately addressed. This work evaluates current RNA simulation force fields concerning Mg²⁺-phosphate and Mg²⁺-water exchanges and investigates the thermodynamics of chelated phosphate-Mg²⁺ coordination, aiming to address key gaps in our understanding of different modes of ion-RNA interactions in RNA-ion-atmospheres.

## Computational Methods

### Systems Preparation for Atomistic Explicit Solvent Simulations

#### BWYV PK System

The initial configuration of the BWYV RNA pseudoknot was obtained from the Protein Data Bank (PDB ID: 437D), corresponding to an X-ray crystal structure reported by Rich and coworkers (**Figure 1A**).^28^ The Amber 99 force field^29^ with parmbsc0^30^ and chiOL3 extensions^31^, was employed to generate the atomistic topologies, for both the RNA and ligand GTP. The system includes a chelated Mg²⁺ ion. The RNA pseudoknot was placed in a cubic simulation box with dimensions of 100 Å × 100 Å × 100 Å. Following DMP systems, the TIP3P water model was used as a solvent model.^32^ To neutralize the overall system charge, a suitable number of Mg²⁺, K⁺, and Cl⁻ ions were added, maintaining concentrations of 2 mM [Mg²⁺] and approximately 100 mM [K⁺], thereby simulating a mixed salt ionic environment within the 100 Å × 100 Å × 100 Å box. Detailed information about the amount of ions and water at different compositions is shown in **Table S1**.

**Figure 1:**
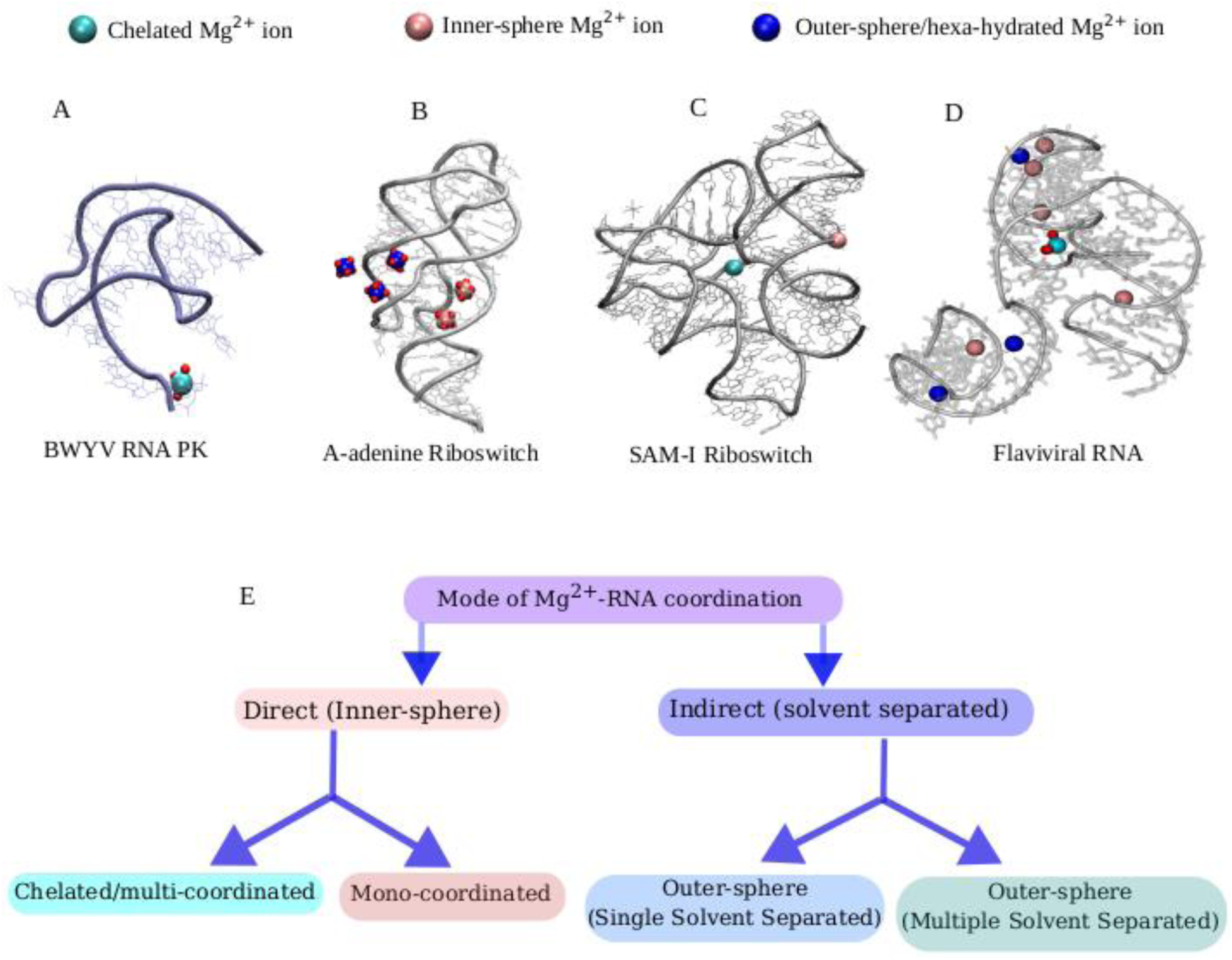
X-Ray Diffraction (XRD) structures of the four different RNA resolved with some Mg^2+^ ion show different modes of Mg^2+^-phosphate interactions (cyan: Chelated, pink: inner-sphere and blue: outer-sphere/hexahydrated Mg^2+^). **(A)** The crystal structure of the Beet Western Yellow Virus (BWYV) shows the presence of chelated/tri-phosphate coordinated Mg^2+^ (in cyan) at the terminal site. Small red beads are co-ordinated water molecule’s oxygen atom associated with the chelated Mg^2+^. **(B)** The structure of the A-adenine riboswitch contains five Mg^2+^, out of these two are mono-phosphate coordinated (pink), and remaining three are outer spheres (blue). Small red beads represent the co-ordinated water molecule’s oxygen atom associated Mg^2+^. **(C)** The structure of S-Adenosyl methionine riboswitch (SAM-I) reveals two type of Mg^2+^: chelated/bi-phosphate coordinated (cyan) and mono-phosphate coordinated (pink). **(D)** Structure of Flaviviral RNA shows the presence of all types of Mg^2+^ interacting with RNA phosphate. **(E)** Modes of Mg^2+^-RNA interactions are classified through a flowchart.

#### A-adenine Riboswicth and SAM-I Riboswitch System

The initial structure of the SAM-I riboswitch bound to its ligand, SAM, was obtained from the Protein Data Bank (PDB ID: 2GIS), as resolved by Batey and co-worker via X-ray crystallography (**Figure 1B**).^33^ Similarly, the crystal structure of the A-adenine riboswitch RNA bound to the ligand ADE was retrieved from PDB ID: 1Y26, as resolved by Serganov et al. (**Figure 1C**).^34^ The ligands SAM and ADE were parameterized for AMBER using quantum mechanical calculations performed with GAMESS ^35^, followed by charge derivation using the Restrained Electrostatic Potential (RESP) method^36^ through the R.E.D. software package.^37^ Additional force field parameters for the ligands were adopted from the Generalized Amber Force Field (GAFF).^38^ The RNA topology for the SAM-I and A-adenine riboswitches was constructed using the same force field parameters applied to the BWYV pseudoknot system.

The crystal structure of the SAM-I riboswitch revealed the presence of two site-specific chelated Mg²⁺ ions, while the A-adenine riboswitch structure contained five RNA-bound Mg²⁺ ions. To neutralize the charges in both systems, an appropriate number of counter ions were added, while retaining the chelated Mg²⁺ ions, and maintaining ionic concentrations of 2 mM [Mg²⁺] (**Table S1**). The same cubic box dimensions and TIP3P water model^32^ used for the BWYV pseudoknot system were employed here as well.

#### Flaviviral RNA System

We have taken an X-ray crystallography structure of Xrn1-resistant Flaviviral RNA from the Murray Valley Encephalitis (MVE) virus (PDB ID: 4PQV), reported by Chapman and co-workers.^39^ The crystal structure revealed 9 site-specific Mg²⁺ ions in the RNA, comprising one chelated ion, five inner-sphere ions, and three outer-sphere ions (**Figure 1D**). Achieving the correct distribution of outer-sphere and inner-sphere ions in the mixed ionic environment (K⁺, Mg²⁺) presented a significant challenge. To neutralize the system’s net charge and replicate physiological conditions with 2 mM [Mg²⁺] and approximately 100 mM [K⁺], appropriate number of counterions were added to the simulation box (**Table S1**). The same box dimensions and water model are employed for the BWYV PK.

#### Mg^2+^-Phosphate Systems

We have created two model systems that resemble the RNA backbone to study magnesium-phosphate interaction and exchange phenomena for inner-sphere mono-phosphate coordinated and chelated/biphosphate coordinated Mg^2+^ ions. Avoiding the topology-specific structural complexity, the inner-sphere mono-phosphate coordinated Mg^2+^ coordination has been studied by taking Mg^2+^-coordinating with a single dimethyl phosphate (DMP) anion. The chelated/ biphosphate coordinated Mg^2+^ has been studied with a complex where one Mg^2+^ interacts with two independent DMP anions. The initial structures of these Mg^2+^-phosphate complexes were taken from the crystal structure of the SAM-I riboswitch (PDB ID: 2GIS).^33^ The Avogadro program was used to add hydrogen atoms to make valency adjustments. AMBER^29^ and CHARMM^40^, two commonly used biomolecular force fields, were employed to parameterize both the DMP systems. Force field parameters for AMBER were obtained from the Generalized Amber Force Field (GAFF)^38^, while for CHARMM, the CHARMM General Force Field (CGenFF)^41^ was utilized within the Swiss Param package.^42^ Both the DMP systems were confined to cubic boxes of dimension 5 Å × 5 Å × 5 Å. An optimized TIP3P water model^32^ was employed, consistent with previous study demonstrating minimal impact of water model variations on Mg^2+^-phosphate kinetic descriptions.^27^ To ensure system neutrality, appropriate counter-ions were added. Mg²⁺-phosphate interactions have been investigated in a physiologically relevant salt concentration of 100 mM [KCl].

### Equilibration Protocols

#### RNA Equilibration

The accurate equilibration of ion distributions around RNA, specifically the outer-sphere and inner-sphere ions, in simulations involving a mixture of counter-ions (K^+^, Mg^2+^) is challenging. This complexity arises from two primary issues: (i) The RNA backbone’s strong electrostatic attraction to Mg^2+^ ions causes rapid condensation of Mg^2+^ onto the RNA without the formation of a proper hydration shell; (ii) Once Mg^2+^ ions interact with the RNA, they may adhere to negatively charged sites, with the energy barrier preventing easy unbinding, even if hydrated. The ions in our simulations are a combination of excess ions which balance the RNA charge and bulk ions accounting physiological ion concentration range. To avoid premature condensation of Mg^2+^ ions onto the RNA, the ions were randomly placed inside the simulation box with larger van der Waals radii. Stochastic dynamics were employed for a 10 ns equilibration, with the RNA constrained and a dielectric constant of 80 to mimic water, ensuring convergence of the electrostatic energy. After filling the simulation box with water, we minimized the system’s energy using the steepest descent method^43^ and annealed it to 300 K over 500 ps with positional restraints of 1000 kJ/mol/nm² applied to both Mg^2+^ ions and RNA. Following our early protocol^21,44^, Mg^2+^ ions were first released and equilibrated for 2 ns, then RNA restraints were progressively reduced from 1000 to 100 to 0 kJ/mol/nm² at constant volume, leading to a total 10 ns NVT equilibration. An additional 10 ns of unrestrained equilibration under constant pressure was performed. After these steps, the systems were subjected to a production MD run. The same equilibration procedure was applied to the Mg²⁺-phosphate system. Detailed simulation parameters are provided in the Supporting Information.

Once the equilibrium trajectory was obtained, we performed free energy calculations for Mg²⁺-phosphate binding across different systems by umbrella sampling method.^45,46^ These included Mg²⁺ mono-coordinated with the phosphate group of a single DMP molecule, as well as Mg²⁺ coordinated with phosphate groups from two DMP molecules. Additionally, a two-dimensional free energy landscape was generated based on the Mg²⁺–O_P_ distances using the well-tempered meta-dynamics method. The details of umbrella sampling and meta-dynamics method have been discussed in the Supporting Information.

### Quantification of Interactive Ion Atmosphere around the RNA Systems Using Preferential Interaction Coefficient

RNA is a highly charged polymer, interacts with ions in solution to maintain electrostatic neutrality and creates a local ion atmosphere, which is very different from bulk concentration. To correctly assess the bulk concentration of each salt component from the mixed salt (Mg^2+^+K^+^+Cl^-^) environment of RNA, we first calculated preferential interaction coefficient (𝛤_𝑖_) quantifies the excess of a specific ion species i near RNA molecule.^19^ Experimental methods such as fluorescence and other spectroscopic studies can measure 𝛤_𝑖_. The total charge (Z) of RNA can be expressed as 𝑍 = ∑_𝑖_ 𝑞_𝑖_ 𝛤_𝑖_, where 𝑞_𝑖_ is the charge of each ionic component. The raw concentration of ions ([i]*) is calculated by counting the number of bulk ions N_i_ and water molecules 𝑁_𝐻2𝑂_, beyond 20 Å from any atom of RNA; both are average quantities over time. The concentration is estimated using the following equation: 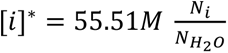, here i is denoted for ionic species such as K^+^, Mg^2+^, and Cl^-^. Although ion densities beyond 20 Å from RNA attain saturation, the charge of RNA cannot be balanced by ions from the computed concentration within it because some charge density is required from out of this limit to balance the residual RNA charge. Hence, the corrected bulk concentrations are calculated by: 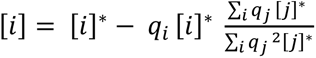, here [i]* or [j]* and 𝑞_𝑖_ are raw concentration and charge, respectively. After calculating the corrected concentrations, the preferential interaction coefficient 𝛤*_i_* is evaluated using the expression: 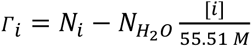. We computed all measurements for the last 100 ns well equilibrated trajectories of the simulations, as shown in **Table S1**, for all four RNA systems, to analyse their salt environment.

## RESULTS AND DISCUSSION

### Generalizing Mg^2+^ Coordination States in the RNA Ion Atmosphere through Atomistic Simulations of Diverse RNA Systems

Recent potential of mean force (PMF) calculations has revealed that K^+^ ions preferentially interact with low electronegative groups such as the oxygen and nitrogen atoms of nucleobases rather than with the phosphate groups in the RNA backbone.^47^ In contrast, inner-sphere Mg^2+^ ions show a strong preference for interacting with RNA’s phosphate groups. Our earlier study also identified such ion-specific preferential interactions of K^+^ and Mg^2+^ in the case of the BWYV pseudoknot (PK).^21^ Atomistic explicit solvent computer simulation methods, particularly those enabling accurate equilibration of ion-atmospheric interaction, are instrumental in distinguishing the microscopic elements of the RNA ion environment. In this study, we have simulated the ion atmospheres of various RNA systems—including the beet western yellows virus (BWYV) pseudoknot, A-adenine riboswitch, SAM-I riboswitch, and Flaviviral RNA pseudoknot —using an all-atom explicit solvent simulation method with a rigorous equilibration pipeline (Equilibrated RMSD trajectories are shown in **Figure S1**). The radial distribution function (RDF) of Mg^2+^ ions around the negatively charged oxygen atoms of the phosphate groups quantitatively reveals distinct peaks corresponding to different Mg^2+^-coordinated complexes, as often observed in crystal structures (**Figure 2**). Based on various direct and indirect interaction between Mg^2+^ and phosphate groups, we have a preliminary classification of the coordination modes as (i) Direct (inner sphere) and (ii) Indirect (solvent separated/outer-sphere) (**Figure 1E**). Direct coordination can further be classified into mono-phosphate coordinated and multi-phosphate coordinated modes. Similarly, solvents-separated interactions can also be classified into single solvent separated and multi-solvent separated, outer-spheric coordination modes. In the equilibrated RNA structures inner-sphere Mg^2+^ ions are found to form one or more coordinated bonds with the oxygen atoms of phosphate groups, as shown in **Figure 1**. Due to the proximity and strong coordination, such Mg^2+^ coordination complexes exhibit a sharp peak at an Mg^2+^-O_P_ distance of ∼1.9–2.0 Å. Additionally, a distinct peak is observed around 4.2 Å, corresponding to an outer-sphere ion. The representative structure corresponding to this peak indicates that the state is single solvent-separated, or hexa-hydrated, Mg^2+^ ion indirectly interacting with the backbone phosphate. This first solvent-separated peak is followed by other solvent-separated minima. An intriguing observation is that as the RNA system complexity increases—from the simple triplex BWYV pseudoknot to the more complex globular RNA like the SAM-I riboswitch—the RDF peaks become more populated. This raises an important question previously posed by Draper and colleagues: could the electrostatic energy contributed by solvent-separated layers (termed “diffuse ions” by Draper) be comparable to the energetic contribution of inner-sphere or chelated ions in RNA structure stabilization.^6^ To fully understand the energetic contributions to RNA structure stabilization, it is crucial to calculate the free energy of RNA-Mg^2+^ binding, taking into account Mg^2+^ ions in both inner-sphere and outer-sphere (diffuse) states. It is worth mentioning here that we are quite aware of the fact that individual RNA-ion interaction may underestimate the collective energetic contribution towards RNA stabilization. Yet, individual ion-RNA interaction free energy calculations provide a basic quantitative estimation and comparative understand keeping aside case-specific topological complexity, which one may add on in the follow up to understand their cooperative/collective effect.

**Figure 2:**
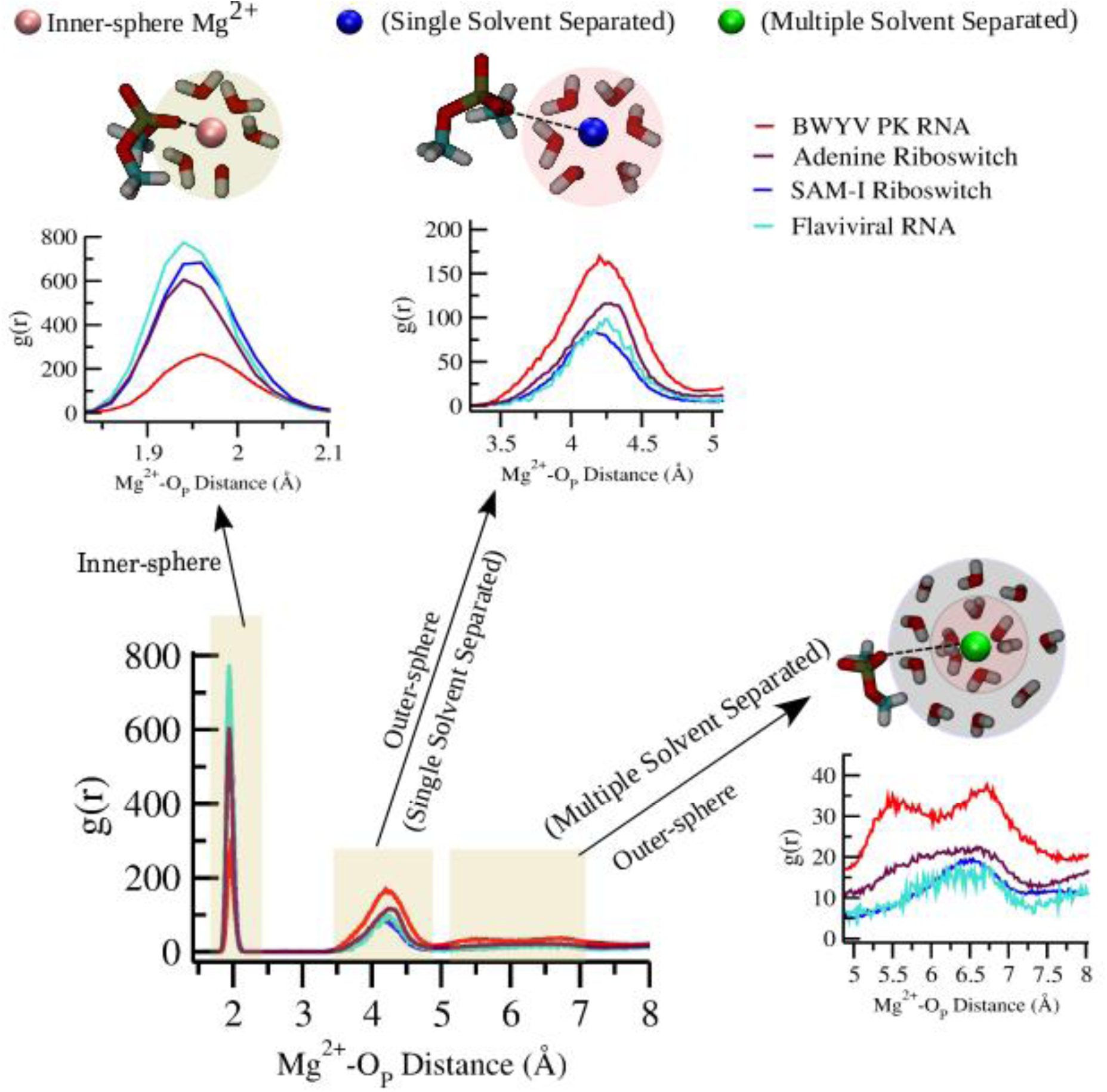
Radial Distribution Function (RDF) calculations to assess different modes of Mg^2+^-RNA coordination in BWYV PK RNA, A-adenine riboswitch, SAM-I riboswitch, and Flaviviral RNA. RDF is plotted as a function of Mg^2+^-O_P_ (oxygen of phosphate) distance. The radial distribution function shows three different modes of interaction namely: Inner-sphere Mg^2+^/Direct coordinated, single-solvent separated outer-sphere and multi-solvent separated (also designated as outer-sphere) Mg^2+^.

### Thermodynamics of Mg^2+^-Phosphate Interaction for Mono-Coordinated Inner-Sphere Complex: Addressing Activation Barrier for Inner-Outer Sphere Ion Exchange and Distance of Closest Approach

Umbrella sampling is a well-established method for quantitatively estimating the free energy difference between the inner- and outer-sphere states of Mg²⁺ interacting with phosphate groups. This approach has been applied to investigate both mono-phosphate-coordinated and bi-phosphate-coordinated (chelated) inner-sphere Mg²⁺ complexes. To characterize the mono-coordinated state, a simplified model system comprising a Mg²⁺ ion and dimethyl phosphate (DMP) was used to avoid the topological complexities inherent in RNA structures. The binding free energy between the phosphate oxygen (O_P_) of DMP and the Mg²⁺ ion was calculated using various Mg²⁺ force-field parameters (**Figure 3**). The corresponding parameter sets are listed in **Table S2**.

**Figure 3.**
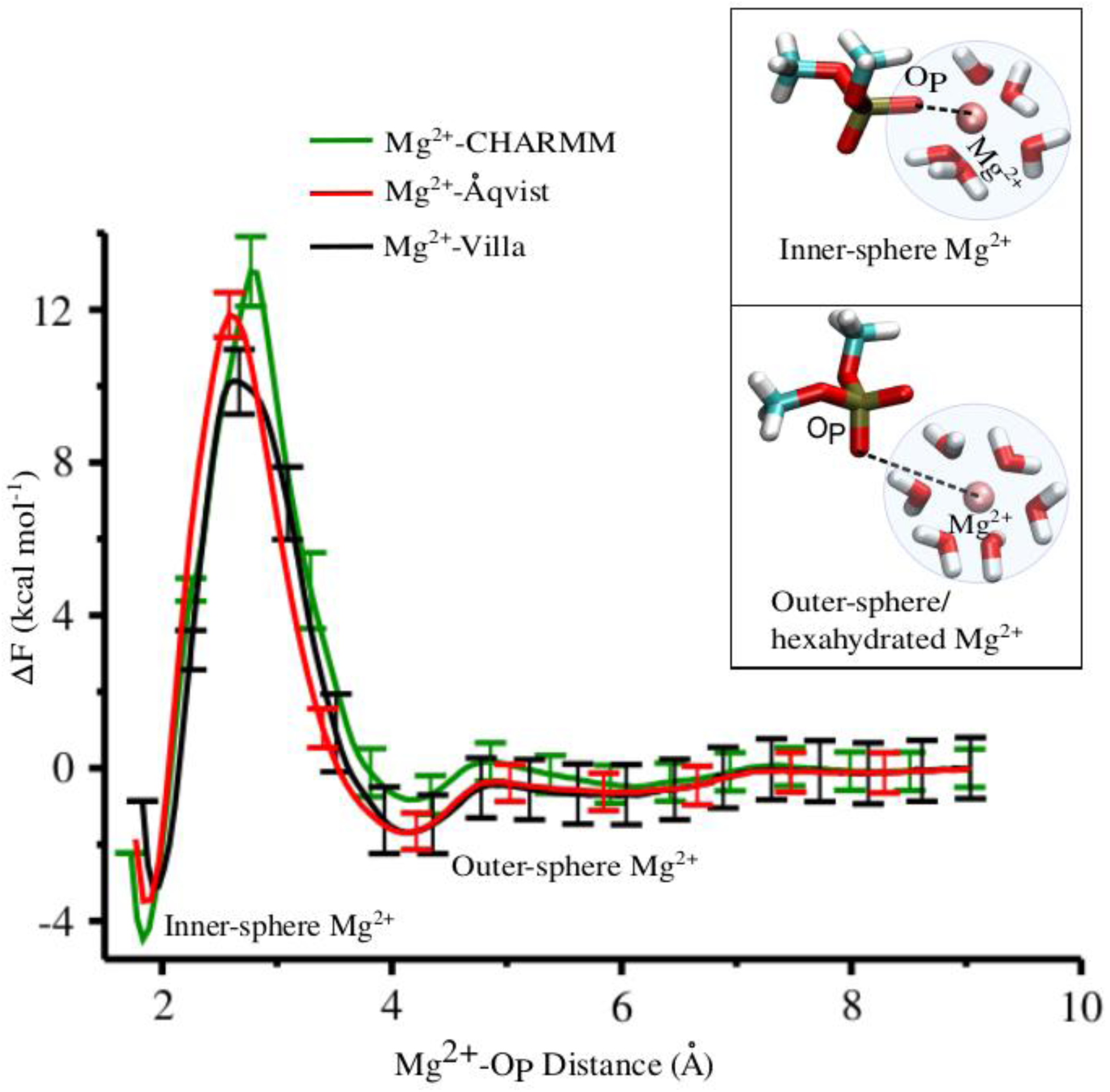
Free energy profiles of inner-sphere Mg^2+^ using different Mg^2+^ parameters. Free energy is calculated as a function of Mg^2+^-O_P_ (oxygen of phosphate) distance using umbrella sampling data. Inset figures show the structure of the system at the two different minima (inner-sphere and outer-sphere states of Mg^2+^).

The free energy plot also reveals a distinct, high-barrier separation between the inner- and outer-sphere Mg²⁺ complexes (**Figure 3**). The barrier estimation is given in **Table S3**. The outer-sphere Mg²⁺ complex, a hexa-hydrated ion, is significantly more stable compared to bulk or diffused hexa-hydrated Mg²⁺. Comparing transition barriers between inner- and outer-sphere states across different Mg²⁺ parameters, earlier work by Villa and co-workers ^27^ identified that Mg^2+^-LB-Åqvist (inner-outer exchange rate: 1.3 × 10⁻³ s⁻¹, water model: TIP3P) and Mg^2+^-CHARMM (inner-outer exchange rate: 2.6 × 10⁻³ s⁻¹, water model: TIP3P) displaying similar slow exchange rates, while the Mg²⁺-Villa parameter exhibited a much faster exchange rate of 10.3 s⁻¹. Although experimental data for phosphate–Mg²⁺ exchange in simple monophosphate systems like DMP are unavailable, existing ²⁵Mg NMR studies for Mg²⁺ interactions with nucleic acids report binding rates of 0.5 × 10³ s⁻¹ for DNA^48^, 1.5 × 10³ s⁻¹ for 5S rRNA^49^, and 2.5 × 10³ s⁻¹ for tRNAᵖʰᵉ ^49^, corresponding to barrier heights of 12.7–13.3 kcal/mol.^27^ While the exchange rates and barrier heights depend on the RNA’s structural topology, the Mg²⁺-Villa parameter consistently provides the closest match to experimental data.

In addition, X-ray diffraction data from a solution of Mg(H₂PO₄)₂ revealed a characteristic Mg–O distance of ∼2.1 Å. Similarly, the free energy plot (**Figure 3**) indicated that the Mg²⁺–O_P_ minimum distance for the inner-sphere mono-phosphate-coordinated Mg²⁺ is ∼2 Å.^50^ A comparison of three Mg²⁺ force-field parameters showed slight variations in the Mg²⁺–O_P_ distance, with the Mg²⁺-Villa parameter closely matching the experimentally determined distance. Additionally, Mg²⁺-Villa parameters closely align with the experimental Mg²⁺–P distance (3.41 Å vs. 3.6 Å reported by Caminiti *et al*. ^50^), outperforming other models, which yielded distances between 3.30 and 3.38 Å (**Figure 4A, 4B, 4C**). Interestingly, the Mg²⁺–O_W_ distance (oxygen of water) followed a similar trend to Mg²⁺–O_P_ (**Figures 4D, 4E, 4F**). Experimentally, distinguishing between these distances is challenging, often resulting in a single peak around ∼2.1 Å in diffraction data. This characterization is crucial because such pairwise minimal approach distances are frequently employed in mean-field and coarse-grained potential development.^51–54^

**Figure 4:**
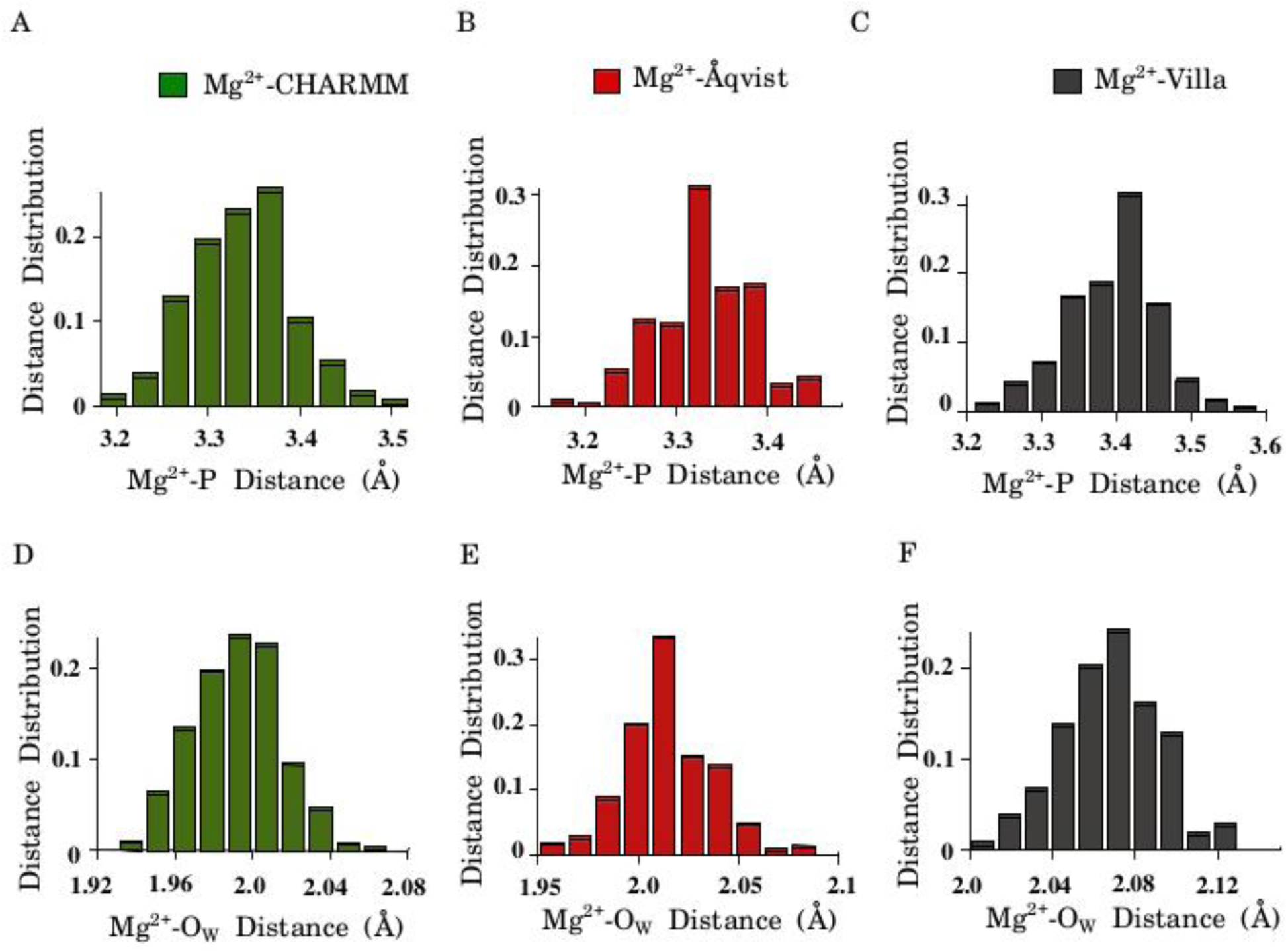
Characteristics distance distribution plots of inner-sphere Mg^2+^ for different Mg^2+^ parameters. **(A)** Distance distribution of Mg^2+^-P for CHARMM parameter. **(B)** Distance distribution of Mg^2+^-P for Åqvist parameter. **(C)** Distance distribution of Mg^2+^-P for Villa parameter. Similarly, the distance distributions of Mg^2+^-O_W_ (oxygen of water) are shown in **(D)**, **(E)**, **(F)**. The most probable values of distances (for Mg^2+^-P and Mg^2+^-O_W_) are greater for Mg^2+^-Villa parameter compared to other Mg^2+^ parameters.

In earlier work, Villa and co-workers used the CHARMM27 force field to model the DMP molecule. Comparatively, newer versions like CHARMM36^40,55^ and AMBER (amber99_parmbc0_chioL3_po with modified dihedral parameters^29,31,56^, reduce the transition barrier between inner- and outer-sphere states and enhance the stability of the outer-sphere complex (**Figure S2**). These updated parameters consistently indicate that the formation of an inner-sphere Mg²⁺–O_P_ bond, involving the breaking of a Mg²⁺–O_W_ bond, is a high-barrier process, but the free energy difference between the states is relatively small. In fact, recent studies on ATP tri-phosphate- and GTP bi-phosphate-coordinated Mg²⁺ complexes^57^ revealed that while the energy barrier between inner- and outer-sphere states is higher for these systems, the thermodynamic stability of both chelated (inner-sphere) and hexa-hydrated (outer-sphere) forms of Mg²⁺ remains comparable. This underscores the equivalence and importance of these Mg²⁺ coordination states as integral components of the RNA ion atmosphere, offering valuable insights for both RNA experimentalists and modelers.

### Free energy Landscape of Mg^2+^ Chelation: Emergence of Dynamic Pre-chelate Complexes Stabilized by Meta-sphere Mg^2+^-Phosphate Coordination

To investigate bi-phosphate-coordinated/chelated Mg^2+^ complex, a simplified model system (representing multi-phosphate ion chelation environment as in RNA) consisting of a Mg^2+^ ion chelated with two dimethyl phosphate (DMP) molecules have been used. The first step involved deriving a two-dimensional free energy landscape based on the Mg²⁺–O_P_ distances (r₁ = Mg²⁺–O_P1_; r₂ = Mg²⁺–O_P2_) using the well-tempered metadynamics method (detailed in the Supporting

Information, **Figure 5A**). A key objective of this study is to assess different coordinated states of Mg²⁺, particularly focusing on the characteristic Mg²⁺–O_P_ distance of their closest approach, which is critical for correlating with the radial distribution function observed in complex RNA systems (**Figure 2**). Therefore, our order parameters are specifically based on distances, implicitly accounting for water-exchange phenomena. The free energy landscape reveals four distinct states: (i) Chelated State: A deep well representing the bi-coordinated (chelated) Mg²⁺ state (**Figure 5B**) at around r_1_≈r_2_≈2.0 Å. (ii) Pre-Chelate State 1 (PC1): If the Mg²⁺–O_P_ distance increases along either r₁ or r₂, a water molecule enters the ion-solvation shell, forming a hexa-coordinated Mg²⁺ complex, while the remaining Mg²⁺–O_P_ distance remains intact, indicating a mono-coordinated state. This species differs from the mono-coordinated species in **Figure 3**, as it involves an outer-sphere phosphate rather than water. As this state may behave either as inner-sphere or outer-sphere complex depending on phosphate reference point before the formation of a chelated form, we label it as the first pre-chelate state (PC1 in **Figure 5C**), where a phosphate group occupies its 1^st^ ion-solvation layer. (iii) Pre-Chelate State 2 (PC2): Further increases in the Mg²⁺–O_P_ distance along r₁ (or r_₂_) lead to a shallow minimum representing the second ion-solvent-separated pre-chelate state (PC2 in **Figure 5D**). (iv) Outer-Sphere Complex: Upon continuing the distance increase along either Mg²⁺–O_P_₁ (r₁) or Mg²⁺–O_P_₂ (r₂), both pathways converge to form a pure outer-sphere complex, surrounded by six water molecules. Three key findings emerge from this analysis are as follows (i) The radial distribution function (RDF) around the same characteristic distance can mix energetically distinct ion species, with their individual contributions to RNA stabilization cannot be distinguished. For instance, energetically a bit different ion-species -chelated and mono-coordinated/prechelate inner-sphere Mg^2+^ share same peak position around Mg^2+^-O_P_ distance∼2.0Å in the RDF as shown in **Figure 2**. These energy states have been distinguished in this 2D free energy analysis. (ii) The free energy difference between the chelated and mono-coordinated states (as observed in **Figure 3**) is negligible, indicating that chelation is not much free-energetically favoured process, compared to the formation of 1^st^ pre-chelated complex, PC1. (iii) It is interesting to note that, in the PC1 and PC2 states, one phosphate group is maintaining a mono-coordination with the central Mg^2+^, while other phosphates are interacting with the central Mg^2+^, in an outer-sphere/solvent manner. Because of such hybrid inner and outer-sphere nature, we refer such type of Mg^2+^-phosphate coordination as “meta-sphere coordination”. (iv) The transition barrier from the chelated state to the mono-coordinated PC1 state was further investigated using umbrella sampling, incorporating ion-coordinated states derived from the metadynamics simulations (**Figure S3**).

**Figure 5.**
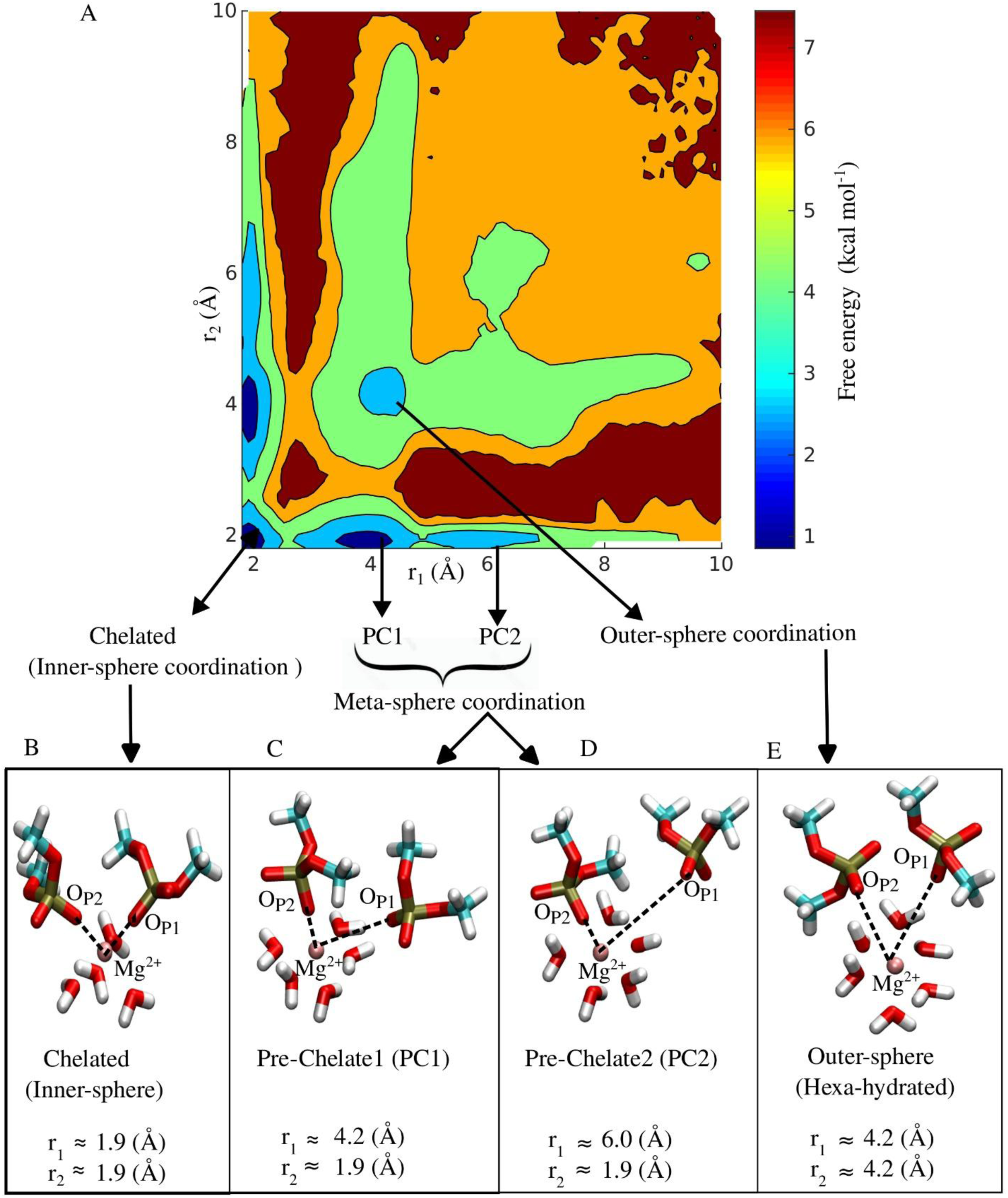
Mechanism of Mg^2+^ chelation. **(A)** Two dimensional free energy landscape of Mg^2+^ is plotted as a function of the distances r_1_ (Mg^2+^-O_P1_) and r_2_ (Mg^2+^-O_P2_) using sampled trajectory from well-tempered metadynamics simulation. The free energy plot majorly finds four stable minima Chelated state (Inner-sphere coordination of Mg^2+^), Pre-Chelate 1 (PC1) and Pre-Chelate 2 (PC2) which are Meta-sphere coordination of Mg^2+^, and the Outer sphere state. (**B)**, (**C)**, (**D)**, (**E**) The representative structures of corresponding minima are shown.

Four distinct states—chelated, two solvent-separated pre-chelate states (PC1 and PC2), and the outer-sphere state—were further analyzed to re-evaluate their stability through equilibrium MD simulations. The trajectories from these simulations provided quantification of the characteristic distances of closest approach for Mg²⁺–P (**Figure S4**) and Mg²⁺–O_W_ (**Figures S5, S6**), for 1^st^ and 2^nd^ ion-solvent-separated layers.

### Dynamic Transitions between Solvent-Separated Pre-Chelate Complex States of Mg²⁺

The most intriguing finding of this study is that the transition between the chelated and outer-sphere states is mediated by the formation of a dynamic ensemble of pre-chelate complexes, which includes both the solvent-separated pre-chelated complexes of Mg²⁺-PC1 and PC2 states. After identifying these states from the metadynamics free energy landscape, we performed classical equilibrium simulations considering both the symmetric type interaction of Mg^2+^ with the two non-local oxygens of phosphate groups, along the r₁ and r₂ directions (**Figures 6A-6F**). From our equilibrium simulation trajectories, we have observed a dynamic exchange between the PC1 (first solvent-separated state) and PC2 (second solvent-separated state), r_1_ and r_2_. In both cases, we observe frequent transitions between the PC1 and PC2 states (**Figures 6E, 6F**), with the probability distribution revealing a larger population of the PC1 state compared to PC2 (**Figures 6G, 6H**). This observation is consistent with the free energy difference derived from the metadynamics calculations (**Figure 5A**). Due to the continuous hopping between the PC1 and PC2 states, we refer to PC1 and PC2 states together as the dynamic pre-chelate ensemble of Mg²⁺.

**Figure 6:**
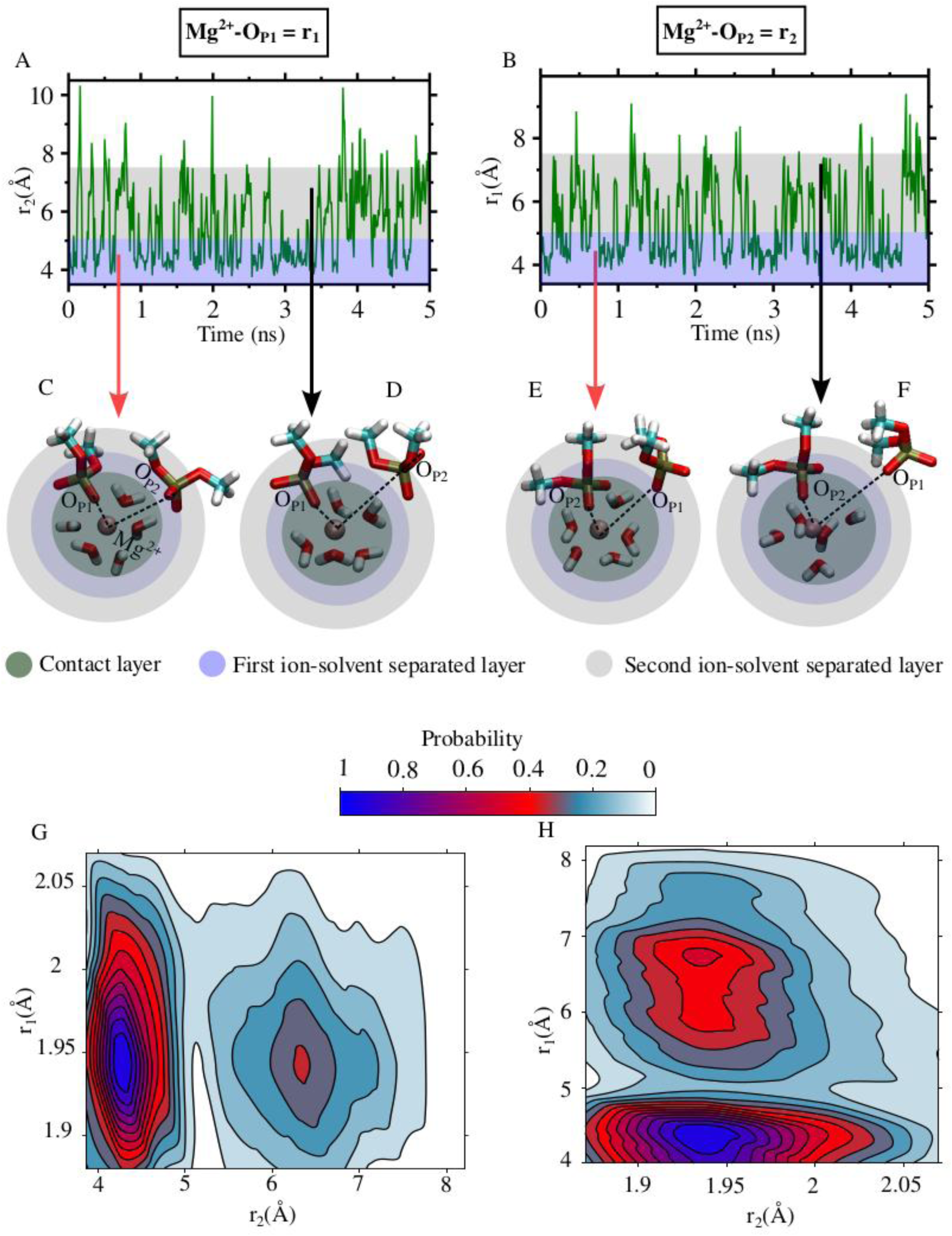
Dynamical transitions of solvent separated phosphate between Pre-Chelate 1 (PC1) and Pre-Chelate 2 (PC2) states. **(A)** Time evolution of distance r_2_ (Mg^2+^-O_P2_). During this analysis, the distance Mg^2+^-O_P1_ or r_1_ (distance for the phosphate which is made direct contact with Mg^2+^) is nearly constant. The blue shaded rectangle represents the 1^st^ ion solvation layer (3.5-5 Å) and the grey shaded region is for 2^nd^ ion solvation layer (5-7.5 Å). **(B)** In a similar way, distance r_1_ is plotted as a function of time while the distance r_2_ is nearly constant. **(C)**, **(D)** The representative structures for 1^st^ and 2^nd^ solvation layer of Mg^2+^ (of Figure **A**) are extracted from the simulation trajectory. The structures show the dynamical hopping between 1^st^ and 2^nd^ solvation layers (essentially between PC1 and PC2 state of Mg^2+^). Similarly **(E)**, **(F)** represents the representative strictures of the solvation layer of Figure **B**. **(G)** 2D contour plot of probabilistic population distribution is represented as function of distances r_1_ and r_2_. This shows that r_1_ is nearly fixed but r_2_ finds two most probable peaks at 4.2 Å (PC1 state) and 6.5 Å (PC2 state). **(H)** Similar analysis like **G**, but in this case r_2_ is nearly constant and r_1_ shows hopping between PC1 and PC2 states.

### Dangling Behaviour of Phosphates in the Solvent Separated Layers of Mg^2+^

From our early analyses we find that the microscopic origin of the emergence of such stable, pre-chelate complex states, PC1 and PC2, underlies in the fluctuation of its ion solvation layers. We suspected a noticeable fluctuation may originate from the reorganization of two local oxygen of an independent solvent separated phosphate group. Nevertheless, the RDF, calculated from the local oxygen atoms, O_P2_ (O_P2_ or O_P2_’ of, local oxygen of phosphate), highlights the peak position a stable first solvent-separated minimum and a broad distribution of low-barrier, solvent-separated states (**Figures 7A**). From equilibrium simulations, we analysed the fluctuations in the distance between these local oxygens of this unbound solvent separated phosphate and the central Mg²⁺ and projected along Mg^2+^-O_P2_ (r_2_) and Mg^2+^-O_P2_’ (r_2_’) presenting it into a 2D probabilistic population distribution plot (**Figure 7B**). Interestingly, this population distribution reveals that two local oxygens (O_P2_ and O_P2’_) of the phosphate ions exhibit a “dangling” behavior, where they competitively interact with the central Mg²⁺. The Mg²⁺–O_P_ RDF, considering any local oxygen of the phosphate, shows equal probability for both oxygens. In fact, the angle distribution of the two local O_P_ vectors relative to the central Mg²⁺ demonstrates the symmetric nature of the phosphate group’s local oxygens (**Figure S7**). Due to this symmetric nature of local oxygens, the transition from PC1 to PC2 involves four dynamic states: (i) **PC1** (r_2_ ≈ r₂’, both local oxygens maintaining the characteristic first solvent-separated distance of ∼4.2 Å), (ii) **IPC1** (r_2_ < r₂’), (iii) **IPC2** (r₂ >r_2_’), and (iv) **PC2** (r_2_≈r_2_’), both local oxygens maintaining the characteristic second solvent-separated distance of ∼6.0 Å). The above phenomena holds true for any solvent-separated phosphate.

**Figure 7:**
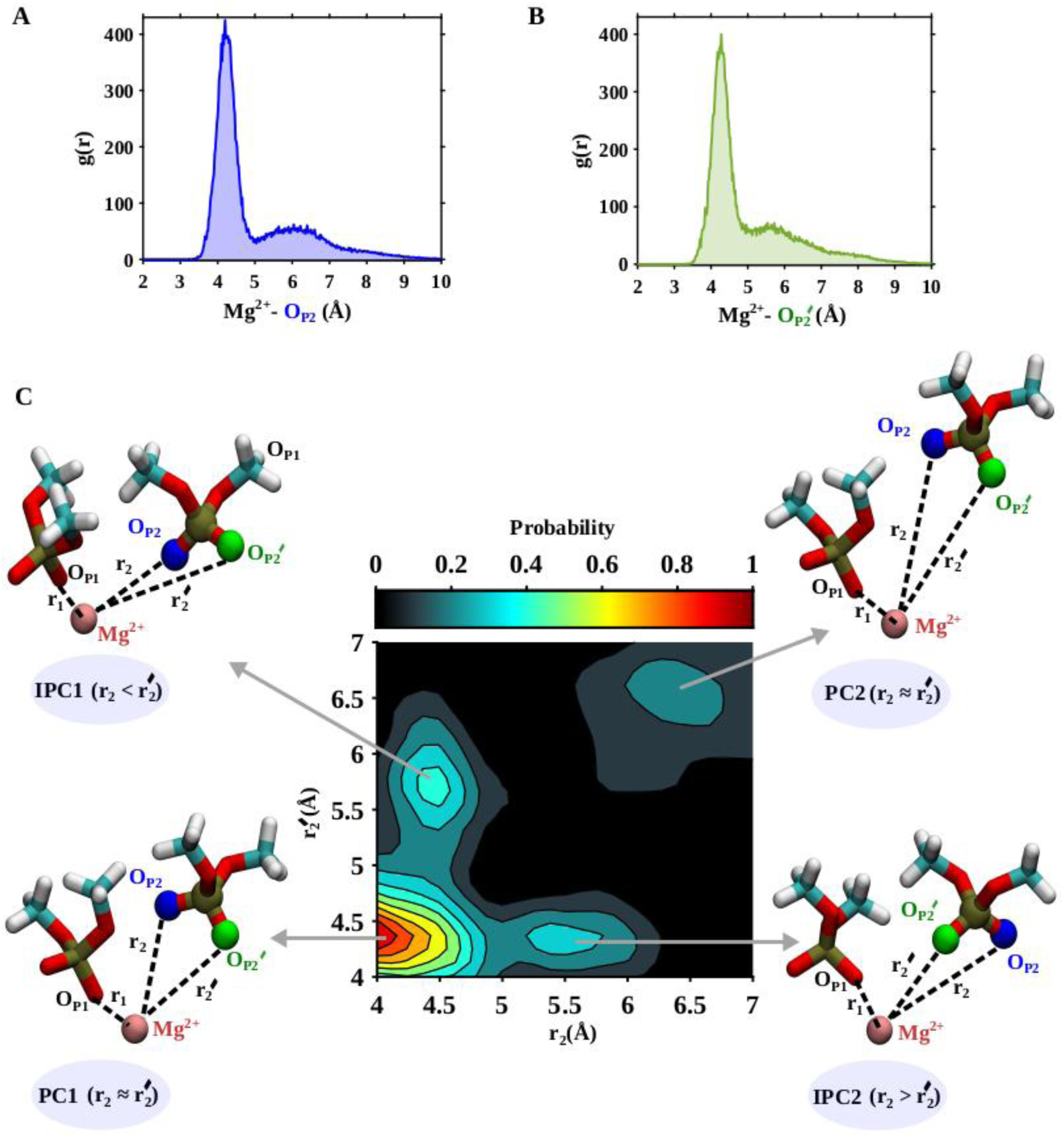
Distance dependent conformations of the solvent separated phosphate relative to Mg^2+^. **(A)** Radial distribution function (g(r)) of the first phosphate oxygen O_P2_ (represented in blue) is plotted with respect to Mg^2+^-O_P2_ distance. **(B)** Radial distribution function (g(r)) of the second phosphate oxygen O_P2’_ (represented in green) is plotted with respect to Mg^2+^-O_P2’_ distance. **(C)** Two-dimensional contour plot of probabilistic population distribution is represented as function of r_2_ and r_2’_, where r_2_, r_2’_ are the distances of O_P2_ and O_P2’_ from Mg^2+^. The plot finds majorly four states (PC1, IPC1, IPC2 and PC2) with significant probability values. The representative structures of the states are shown.

### Oxygen Exchange Mechanism between Phosphate and Water Ligands in Ion-Solvent-Separated Layers of Mg^2+^

The calculation of the average number of water ligands around Mg²⁺ revealed that during the transition paths (PC1 → IPC1 → PC2 or PC1 → IPC2 → PC2), water molecules are incorporated into the first and second solvation layers, while the contact layer of Mg²⁺ remains intact (**Figure 8A**). To further understand this water inclusion and phosphate reorganization, we quantified the number of bridged water molecules connecting Mg²⁺ to the local oxygen atoms of the phosphate group (**Figure 8B**), using a graph-theoretical approach (described in the Methods section) to identify the shortest path length. In the PC1 state, both local phosphate oxygens are part of the first ion solvation layer of Mg²⁺, forming hydrogen bonds with directly coordinated water molecules (**Figure 8C**). In the IPC1 and IPC2 states, representative structures show clear evidence of oxygen exchange from phosphate to water. The dangling behavior of the relatively free (outer-sphere) phosphate oxygens facilitates the inclusion of additional water molecules to replenish the adjacent solvation layer. In the PC2 state, the number of bridged water molecules and the corresponding structures indicate the inclusion of two additional water molecules, displacing both phosphate oxygens from the first to the second solvation layer.

**Figure 8.**
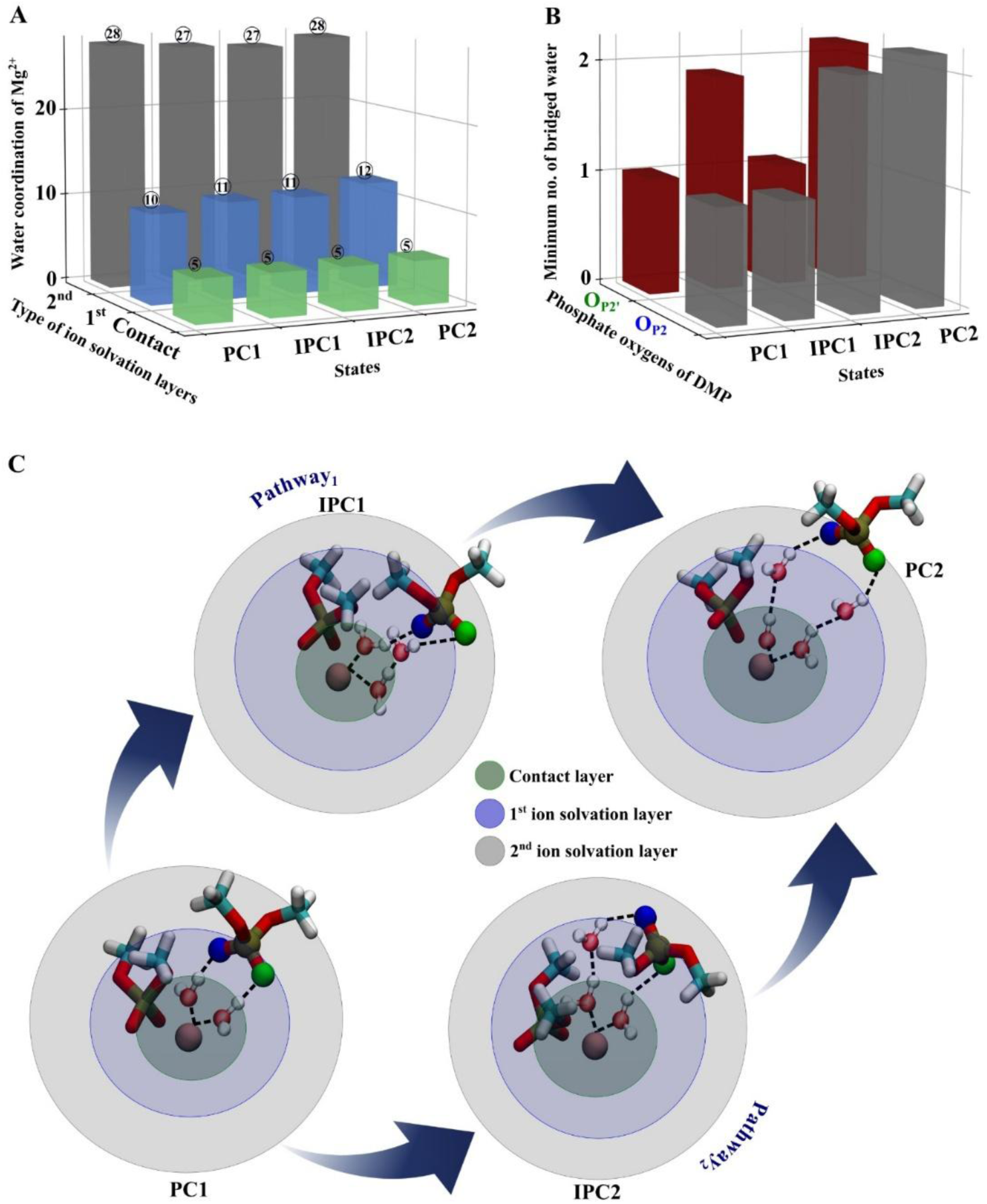
Bridged water molecules between the Mg^2+^ and oxygens (O_P2_/O_P2’_) of the solvent separated phosphate. (A) Average number of co-ordinated water molecules in the different layers of Mg^2+^ (contact layer, 1^st^ ion solvation and 2^nd^ ion solvation layers) for all the states (PC1, IPC1, IPC2 and PC2) are represented as a 3D transparent bar plot. Three distinguished colours are used to represent different solvation layers of Mg^2+^ (contact layer: light green; 1^st^ ion solvation layer: cornflower blue and 2^nd^ ion solvation layer: grey). (B) Minimum no. of bridged water molecules between Mg^2+^ and phosphate oxygens for all the states (PC1, IPC1, IPC2 and PC2) are represented as a 3D transparent bar plot. The first phosphate oxygen O_P2_ is shown in blue and the corresponding bridged water no. to this interaction site is shown in grey bar; the second phosphate oxygen O_P2’_ is shown in green and the bridged water no. to this interaction site is shown in red bar. (C) The bridged water molecules from the Mg^2+^ to the specific type of oxygens of the solvent separated phosphate (O_P2_/ O_P2’_) for the states PC1, IPC1, IPC2 and PC2 are shown in black dotted lines. There are two probable pathways by which PC1 state goes to PC2 state (Pathway_1_: PC1 → IPC1 → PC2 and Pathway_2_: PC1 → IPC2 → PC2). The pathways are shown in curved blue arrows. Only the bridged water molecules (not all the co-ordinated water molecules) are shown inside the solvation layers of Mg^2+^. The representative structures are extracted from the simulation trajectories of the individual states.

### Mg^2+^ Chelation Mechanism in Complex RNA Structure: Validating the Emergence Pre-chelate States in the Tertiary Phosphate Network of RNA

While our earlier explorations were based on a simple model system of Mg²⁺ chelated with two phosphate groups from two DMP molecules, the emergence of the pre-chelate complex state of Mg²⁺ underscores the critical role of unbound phosphate ligands in its solvent-separated ion layer. To further validate the formation and stability of such pre-chelate complexes, we examined the binding free energy profiles (calculation method is discussed as Supporting Information) of a chelated Mg²⁺ ions in an RNA structure of SAM-1 RNA aptamer.^33^ This Mg^2+^ is located at the four-way junction which is the central core of this RNA, surrounded by multiple adjacent phosphate groups. This core-chelated Mg^2+^ bridges two bases A10 and U64 (**Figure 9A**). The binding free energy is calculated as function of a distance between chelated Mg^2+^ and the center of mass of O_p_ of A10 and O_p_ of U64 (**Figure 9B**). Like earlier, in this free energy profile of chelated Mg^2+^, we obtain five distinct wells corresponding to five barrier-separated state: (i) Chelated State (C) (ii) First solvent-separated meta-sphere coordinated pre-chelate complex (PC1), (iii) Second solvent-separated meta-sphere coordinated pre-chelate complex (PC2), (iv) a broad distribution of another meta-sphere coordinated pre-chelate complex (PC3) (Representative structure has been shown in **Figure S8**) , and (v) a hexa-hydrated outer-sphere complex. The PC1 state, a stable meta-sphere-coordinated state, exhibited a calculated Mg²⁺–O_P_ mean distance of ∼1.95 Å, close to the experimental value of 2.1 Å. Similarly, the Mg²⁺–P mean distance in PC1 is calculated as ∼3.4 Å (experimental value ∼3.6 Å) (**Figure S9**). While PC1, PC2 and PC3, all represent meta-sphere phosphate coordinated pre-chelate state, here we only demonstrate the analysis of PC1. From the atomistic equilibrium simulations of PC1 state we show the dangling motion of local oxygens of two flexible solvent-separated phosphate groups. However, unlike early simple model system, where two local oxygens of flexible phosphate showed symmetric nature and similar electrostatic attraction towards central Mg^2+^, for RNA structure, we do not find the two local oxygen to behave equally. Here, among two local oxygens, which is topologically more closer to the chelated Mg^2+^ (O_P2’_), show less switching between 1^st^ and 2^nd^ ion-solvation layers than the other oxygen (O_P2_), attributing to the dynamic transition between PC1 and PC2 states (**Figure 9C-9F**).

**Figure 9.**
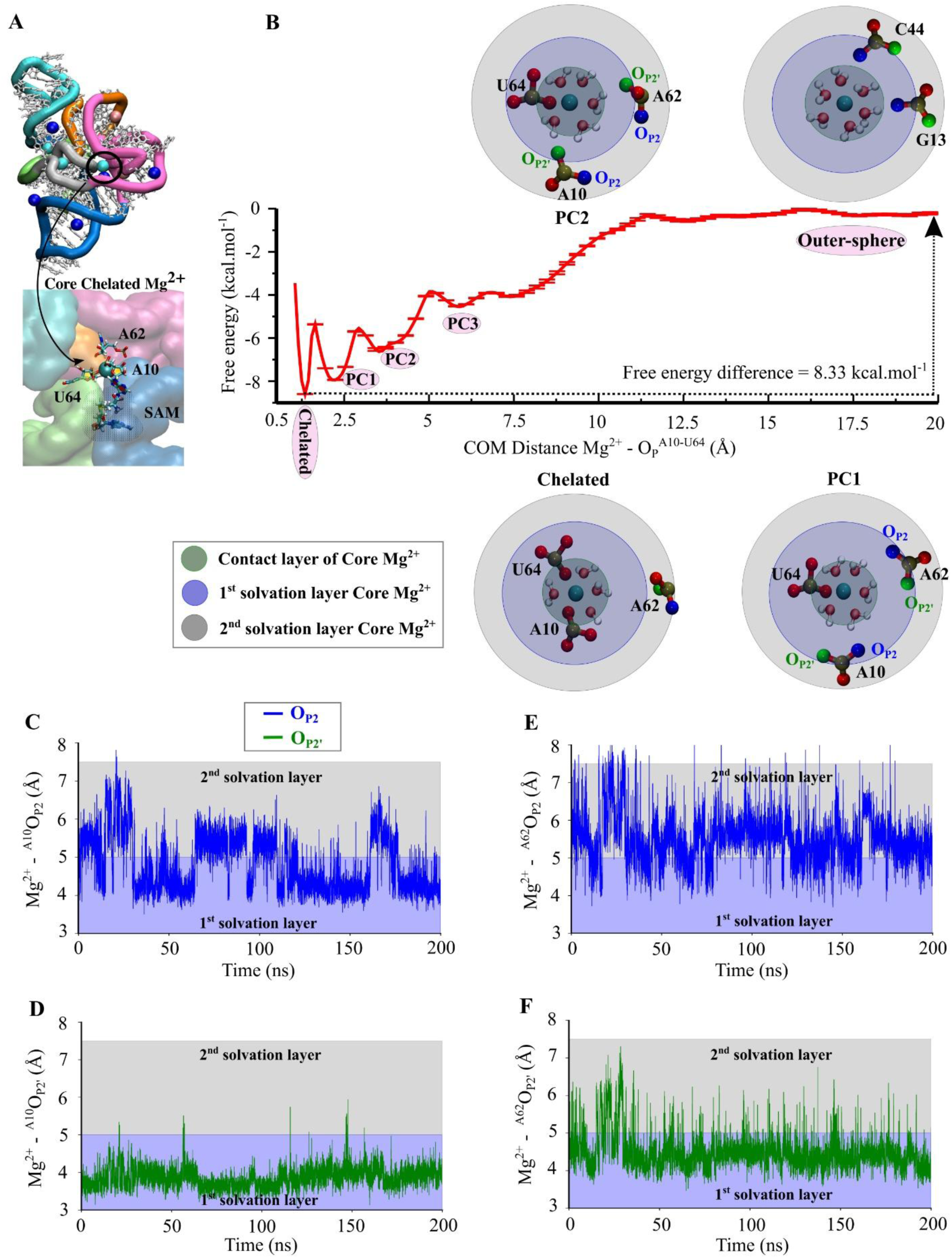
Free energy calculations of the core chelated Mg^2+^ of SAM-I riboswitch RNA in multi-phosphate environment and characteristics behaviour of the solvent separated phosphates. **(A)** Equilibrium structure of SAM-I RNA along with its interactive Mg^2+^ ion environment. Mg^2+^ ions are represented in three distinguished colours (cyan: chelated; pink: single phosphate co-ordinated inner sphere; blue: outer sphere) in van der waals bead representation. The interacting RNA residues of core chelated Mg^2+^ are shown (core Mg^2+^ is chelated by the direct contact of A10 and U64; A62 remains as multi-solvent separated outer sphere phosphate) in the zoomed image. Ligand (SAM) binding pocket is located beneath the core Mg^2+^. The snapshot is extracted from one of the equilibrium simulation trajectories of SAM-I maintaining ∼100 mM KCl, ∼2.0 mM bulk Mg^2+^ concentration. **(B)** Free energy profile of the core chelated Mg^2+^ is represented as a function of Center Of Mass (COM) distance between Mg^2+^ and phosphate oxygens (O_P_) of A10, U64. The representative structures for the corresponding minima (Chelated, PC1, PC2 and Outer-sphere) are shown with different solvation layers of core chelated Mg^2+^. There are two phosphate residues (A10 and A62) in the 1^st^ solvation layer of Mg^2+^ at PC1 and PC2 states. The interacting phosphate oxygens of both the residues A10, A62 are shown in two different colours (O_P2_: blue and O_P2’_: green). **(C)**, **(D)** represents the time evolution of Mg^2+^-O_P2_ and Mg^2+^-O_P2’_ distances for residue A10. The distance trajectory is obtained from the equilibrium simulation of the PC1 state. Similarly, **(E)** and **(F)** represents the time evolutions of Mg^2+^-O_P2_ and Mg^2+^-O_P2’_ distances for residue A62.

## Conclusion

This study of the RNA ion atmosphere presents two key sets of conclusions: a set of quantitative insights derived from free energy calculations, offering foundational data for energy modeling, and a mechanistic insight into magnesium chelation, elucidating dynamic behaviour in multi-phosphate tertiary environment critical for understanding RNA-ion interactions. Both are highly relevant to RNA forcefield developers, structure-based model creators, and RNA experimentalists.

### Quantitative Findings from Free Energy Calculations

Atomistic simulations of RNA structures with varying complexity reveal generalized layers of Mg²⁺-phosphate coordination: the directly coordinated inner-sphere and various solvent-separated outer-sphere ions. Radial distribution function (RDF) analysis highlights distinct peaks for these coordination states and defines their minimal approach distances, consistent with solution X-ray diffraction data. Addressing challenges in Mg²⁺-forcefield parameters, this study confirms earlier findings by Villa et al. that modified Mg²⁺ parameters improve the kinetic barrier of Mg²⁺-water exchange.^27^ In this study, we further demonstrate that Villa’s Mg^2+^ parameter and AMBER force-field^29,31,56^ together achieve a significantly improved kinetic barrier for Mg²⁺-phosphate binding (∼12.9 kcal/mol), closely aligning with experimental values (∼12.7–13.3 kcal/mol).^48,49,58^ However, the free energy difference between inner- and outer-sphere Mg²⁺ states is marginal (∼1.0-2.0 kcal/mol). While these calculations align with previous mono-phosphate Mg²⁺ studies, our exploration of Mg²⁺ chelation involving two phosphate groups provides interesting insights into Mg^2+^-phosphate interaction for RNA. The results indicate that while the kinetic barrier for transitioning between mono-coordinated and chelated Mg²⁺ is substantial (∼12 kcal/mol), again, chelation offers limited free-energy advantage compared to mono-coordination. It is also important to note that the kinetic barrier for phosphate–Mg²⁺ exchange can vary depending on the specific system under study. Variations in system complexity alter local phosphate coordination and the solvation environment around Mg²⁺, leading to distinct pathways and activation barriers. For example, the barrier for transitioning from a chelated state to a fully hexa-hydrated outer-sphere state in the SAM-I aptamer RNA is approximately 8.33 kcal/mol (**Figure 9B**).

### Mechanistic Insights into Magnesium Chelation

A critical finding of this study is the role of dynamic pre-chelate complexes in the transition between chelated and outer-sphere Mg²⁺ states. These transitions are facilitated by solvent-separated phosphate groups that exhibit dynamic, symmetric fluctuations. This behavior allows for the identification of four distinct solvent-separated states (PC1, IPC1, IPC2, and PC2) with characteristic mean Mg²⁺-phosphate oxygen distances (∼4.2 Å for PC1, ∼5.5 Å for IPC1/IPC2, and ∼6.0 Å for PC2), explaining the microscopic origin of the solvent-separated layers in the RDF. Transition pathways highlight a unique oxygen exchange mechanism, where additional water molecules replace phosphate oxygens in the first solvation layer, pushing them into the second layer. This dynamic reorganization underscores the critical role of solvent-separated, flexible phosphate groups in modulating Mg²⁺ solvation and coordination. In complex RNA environments, such as the SAM-I RNA aptamer, free energy calculations reveal stable dynamic pre-chelate ensembles. Here, Mg²⁺ is mono-coordinated with one phosphate group, while two additional phosphates from the first ion-solvation shell exert competitive, water-screened electrostatic interactions. This interplay, reflected in RDF peaks, supports the dynamic nature of Mg²⁺ coordination.

Draper and colleagues have emphasized the dominant role of diffuse ions in RNA stabilization, arguing that site-specific Mg²⁺ chelation contributes minimally due to the high dehydration and ion repulsion penalties.^6,59,60^ Expanding on this perspective, our study introduces dynamic pre-chelate complexes as a compensatory mechanism, where solvent-separated, flexible phosphates facilitate meta-sphere ion-phosphate coordination. These outer-sphere phosphates not only retain additional hydration but also help accumulate diffuse Mg²⁺ within their solvation shells. It is important to clarify that this study does not claim that either diffuse ions or site-specific chelation is universally superior for RNA stabilization; their relative contributions are highly system-dependent. In multi-phosphate tertiary environments of RNA, pre-chelate complexes may or may not outcompete the collective stabilization from diffuse or outer-sphere ions, but this study underscores that ion chelation is neither as energetically unfavourable nor as rare as previously assumed.^6,59,60^ Crucially, the existence of meta-sphere coordination bridging inner- and outer-sphere ion coordination, highlights the importance of tertiary phosphate network, stabilizing an RNA-ion interaction. Therefore, models relying solely on discrete ion-binding sites may oversimplify and misrepresent ion-RNA interactions. In addition to meta-sphere ion-phosphate coordination, quantitative characterization of pure inner and outer-sphere modes of ion-phosphate coordination may provide quantitative insights for refining RNA forcefields and structure-based modelling. In future, we aim to explore the structural and functional relevance of pre-chelate complexes by incorporating dynamic exchange between inner- and outer-sphere ion components into broader RNA folding models, advancing our understanding of ion-mediated RNA stabilization mechanisms.

### Supporting Information

Simulation Parameters Details; Free Energy Calculation Using Umbrella Sampling Method: Constant Velocity Steered MD, Potential of Mean Force (PMF) Calculation; Free Energy Simulation Using Well-Tempered Metadynamics; Number of Bridged Water Calculation; Table S1: Characterization of Ion-atmosphere around Different RNA; Table S2: Different Force fields and Lennard-Jones Parameters; Table S3: Estimation of Activation Barrier for Inner-Outer Sphere in Mg^2+^-Phosphate Exchange for Different Mg^2+^ Parameters; Supporting Figures S1 to S9; ReferencesNotes

The authors declare no competing financial interests.

## Supporting information

https://drive.google.com/file/d/1IKDz6nW0IR0VM7Hp67hXk4Yw-O06mWVG/view?usp=drive_link

## Acknowledgements

SR acknowledges support from the Department of Biotechnology (DBT) (Grant No. BT/12/IYBA/2019/12 and BT/PR40192/BTIS/137/692023).

## Notes

### Competing Interest Statement

The authors have declared no competing interest.

