## Supplementary material for "Free Energy Landscape of Magnesium Chelation Reveals Dynamic Pre-Chelate Complexes Stabilized by Meta-Sphere RNA-Ion Coordination": https://drive.google.com/file/d/1IKDz6nW0IR0VM7Hp67hXk4Yw-O06mWVG/view?usp=drive_link

Akhilesh Jaiswar, Raju Sarkar, Avijit Mainan, Rimi Kundu, Susmita Roy*[†]
[†] Department of Chemical Sciences, Indian Institute of Science Education and Research Kolkata, West Bengal 741246

* Corresponding author


A.J. and R.S. contributed equally

**This PDF file includes:**
**Supporting Information:**

- Simulation Parameters Details
- Free Energy Calculation Using Umbrella Sampling Method
  - *Constant Velocity Steered MD*
  - *Potential of Mean Force (PMF) Calculation*
- **Free Energy Simulation Using Well-Tempered Metadynamics**
- **Number of Bridged Water Calculation**
- **Table S1:** Characterization of Ion-atmosphere around Different RNA.
- **Table S2:** Different Force fields and Lennard-Jones Parameters.
- **Table S3:** Estimation of Activation Barrier for Inner-Outer Sphere in $Mg^{2+}$-Phosphate Exchange for Different $Mg^{2+}$ Parameters
- **Figure S1 to S9**
- **References**

### Simulation Parameters Details

All simulations were conducted using the GROMACS 2018.3 software package.[1,2] The Verlet integrator[3], with 2 femtosecond time step, was employed for integrating the equations of motion. Modified $K^+$ ion parameters were utilized to prevent crystallization at elevated concentrations.[4] Long-range electrostatic interactions were calculated using the Ewald algorithm[5], with a grid spacing of 1.2 Å and a Coulomb cutoff distance of 10 Å. Non-bonded interactions were treated with a van der Waals cutoff of 10 Å. Temperature coupling of the systems were maintained with the Nose-Hoover thermostat[6], and covalent bonds involving hydrogen atoms were constrained using the LINCS algorithm.[7] Different sets of $Mg^{2+}$ parameters have been used to investigate the thermodynamics of $Mg^{2+}$-phosphate interaction (**TABLE S2**).

### Free Energy Calculation Using Umbrella Sampling Method

*Constant Velocity Steered MD*

To explore one dimensional free energy profile of mono-phosphate-coordinated $Mg^{2+}$-DMP system, we employed constant velocity steered molecular dynamics (cv-SMD) simulations. This technique systematically applies a time-dependent external force along the reaction coordinate, overcoming the high energy barrier that prevents spontaneous unbinding at ambient temperatures. A harmonic bias potential $w(r) = \frac{1}{2}kr^2$ along the reaction coordinate r facilitates the unbinding event, systematically pulling the $Mg^{2+}$ ion away from the DMP at a constant velocity. GROMACS 2018.3 software was used to perform the cv-SMD simulations, with the reaction coordinate defined as the center of mass distance between $Mg^{2+}$ and the phosphate oxygen ($O_P$) of DMP.  A force constant of k=4000 kJ/mol/nm² and a pulling speed of 0.00001 nm/ps were applied during the simulations.

Similarly, the binding free energy of core chelated $Mg^{2+}$ in SAM-I RNA (**Figure 9A** in main text) was calculated taking distance between chelated $Mg^{2+}$ and the center of mass of $O_p$ of A10 and $O_p$ of U64 (**Figure 9B** in main manuscript) as a reaction coordinate. In this case, we used a force constant of k=1000 kJ/mol/nm² and a pulling speed of 0.00001 nm/ps during cv-SMD simulations.

*Potential of Mean Force (PMF) Calculation*

For the calculation of PMF profiles, we employed the umbrella sampling method.[8,9] The trajectories from cv-SMD simulations along the reaction coordinate were used to generate multiple umbrella sampling windows, spanning from 1.8 Å to 10 Å. Initial configurations for each window were selected asymmetrically to ensure adequate overlap in the defined phase space. Each window was simulated in the presence of harmonic restraint for 3 ns under the NVT ensemble at 300 K. Larger value of force constants were applied near the high energy barrier regions. Temperature was controlled using a Nosé-Hoover thermostat. Overlap between the windows was ensured to provide comprehensive sampling coverage across the entire phase space along the reaction coordinate, as illustrated in **Figure S4**. In case of Core chelated $Mg^{2+}$ in SAM-I RNA, we generated 44 windows along defined reaction coordinate (mentioned above), which is the distance between chelated $Mg^{2+}$ and COM of phosphates ($O_P$) A10 and U64. The initial COM distance was 1.8 Å. We performed 20ns long simulation for each window. The weighted histogram analysis method (WHAM) was subsequently applied to obtain the PMF profiles, resulting in an unbiased free energy profile. When the center of mass (com) distance is constrained, the solute-solute connecting vector can still rotate freely, allowing for sampling of larger volume elements at greater distances. As a result, an entropic term, given by $S = 2k_B T \ln(r)$, contributes to the average constraint force and must be subtracted accordingly. The final corrected PMF was calculated by the following expression, $V^{PMF} = -k_B T \ln P(r) + 2k_B T \ln(r)$, where P(r) is the probability distribution obtained from WHAM. All potential of mean force (PMF) curves have been normalized to a zero value, using the bulk as a reference. The associated error bars were estimated using the bootstrap method implemented in GROMACS.[10]

**Free Energy Simulation Using Well-Tempered Metadynamics**

To investigate the mechanism of $Mg^{2+}$ chelation with a simplified model system consisting of a $Mg^{2+}$ ion chelated with two dimethyl phosphate (DMP) molecules, we employed 545 ns long well-tempered metadynamics simulations[11] using the same simulation parameters mentioned above. This approach was used to map the 2D free energy landscape (**Figure 5** in the main text) as a function of two collective variables (CVs) namely, $r_1$ and $r_2$ (where, $r_1 = Mg^{2+}–O_{P1}$; $r_2 = Mg^{2+}–O_{P2}$ representing the distances between $Mg^{2+}$ and, the $O_{P1}$ and $O_{P2}$ atoms of the two DMP molecules in double DMP-$Mg^{2+}$ complex. For both the variables, we set a sigma of 0.1 Å, a Gaussian height of 0.5 kcal/mol and a bias factor of 12. During the metadynamics run, we used a hill deposition rate of 0.5 ps. After the simulation, a two dimensional free energy lansdcape was mapped by population based Boltzman probasbility distributions.

**Number of Bridged Water Calculation**

After configuring out the distance dependent conformations of solvent separated phosphate with respect to $Mg^{2+}$, we extracted the static structures of PC1, IPC1, IPC2, and PC2 states (as shown in **Figure 7C** in main manuscript). To classify these conformations based on water inclusion connecting $Mg^{2+}$ to the local oxygen atoms of the phosphate group, we performed additional 8 ns of simulations (2 ns per conformation). During these simulations, the two DMP molecules and $Mg^{2+}$ were position-restrained with a force constant of 1000 kJ/mol.nm$^2$, allowing the water to equilibrate while preserving the orientations of the DMP molecules.

To determine the number of water molecules bridging $Mg^{2+}$ and local oxygen atoms ($O_{P2}$ and $O_{P2'}$) of the solvent separated phosphate, we constructed a distance-based graph where $Mg^{2+}$, $O_{P2}$, $O_{P2'}$ and all oxygen atoms ($O_W$) of the related water molecules were defined as nodes. An undirected edge was assigned between any two nodes within 3.5 Å of each other. To avoid the unnecessary computation, only the water molecules lying within 7.5 Å around $Mg^{2+}$ were considered in the graph, as it is the maximum extension of 2$^{nd}$ solvation layer of $Mg^{2+}$ (mentioned in **Figure 6** of main manuscript). To find out the minimum no. of bridged water molecules, the shortest path between $Mg^{2+}$ (as starting node) and oxygen of interest either $O_{P2}$ or $O_{P2'}$ (as end node) is evaluated by Breadth-First Search (BFS) algorithm.[12,13] The shortest path length in this graph indicates how many steps connect $Mg^{2+}$ and the phosphate oxygen. Since BFS yields the total number of edges in that path, the number of bridging waters is calculated as number of edges – 1. For instance, a path $Mg^{2+} \rightarrow W1 \rightarrow O_{P2}/ O_{P2'}$ has two edges, indicating one bridged water (W1); while the path $Mg^{2+} \rightarrow W1 \rightarrow W2 \rightarrow O_{P2}/ O_{P2'}$ has three edges, signifying two bridged water molecules (W1 and W2). We repeated this process for every frame along the 2 ns simulation trajectory and calculated the time averaged value of minimum no. bridged water.

**Table S1**: Characterization of ion atmosphere around different RNA molecules. Raw salt concentration []* and corrected bulk concentrations [] are determined (using the aforesaid approach) and calculated preferential interaction coefficients, Γ.

| **RNA Systems** | **BWYV PK** | **SAM-I Riboswitch** | **A-adenine Riboswitch** | **Flaviviral RNA** |
|---|---|---|---|---|
| $N_{Mg^{2+}}$ | 6 | 11 | 11 | 13 |
| $N_{K^+}$ | 70 | 116 | 96 | 106 |
| $N_{Cl^-}$ | 54 | 46 | 48 | 65 |
| $[Mg^{2+}]^*$ mM$^a$ | 1.64 | 2.8 | 0.64 | 1.74 |
| $[K^+]^*$ mM$^a$ | 98.64 | 112 | 108.3 | 132.67 |

| | | | | |
|---|---|---|---|---|
| $[Cl^-]^*$ mM[a] | 98.91 | 95 | 94.84 | 126.06 |
| $[Mg^{2+}]$ mM[b] | 1.59 | 2.2 | 0.54 | 1.6 |
| $[K^+]$ mM[b] | 97.18 | 100.5 | 100.54 | 127.64 |
| $[Cl^-]$ mM[b] | 100.3 | 105 | 101.63 | 130.83 |
| $\Gamma_{Mg}^{2+}$ | 6.07 | 9.74 | 10.69 | 12.08 |
| $\Gamma_{K}^{+}$ | 13.42 | 58.59 | 38.34 | 32.84 |
| $\Gamma_{Cl}^{-}$ | -4.42 | -13.98 | -10.28 | -9.98 |

a Raw concentrations are determined 20 Å beyond RNA (asterisk),

b Corrected bulk concentrations are determined using a small potential perturbation approximation method.

**Table S2:** Different Force fields and Lennard-Jones Parameters.

| Force Fields | Authors | Sigma (σ) (Å) | Epsilon (ε) (kcal mol$^{-1}$) |
|---|---|---|---|
| CHARMM 27 | Villa et al. | 2.77 | 0.00295 |
| CHARMM 36-jul2021 | CHARMM | 2.11 | 0.01501 |
| amber99_parmbsc0_chiOL3_po | Åqvist et al. | 1.41 | 0.08974 |
| amber99_parmbsc0_chiOL3_po | Villa et al. | 2.77 | 0.00295 |

**Table S3:** Estimation of Activation Barrier for Inner-Outer Sphere in $Mg^{2+}$-Phosphate Exchange for Different $Mg^{2+}$ Parameters

| Force Fields and $Mg^{2+}$ parameter | Activation Barrier (kcal mol$^{-1}$) |
|---|---|
| CHARMM | 17.8 |
| $Mg^{2+}$-Åqvist | 15.4 |
| $Mg^{2+}$-Villa | 12.9 |

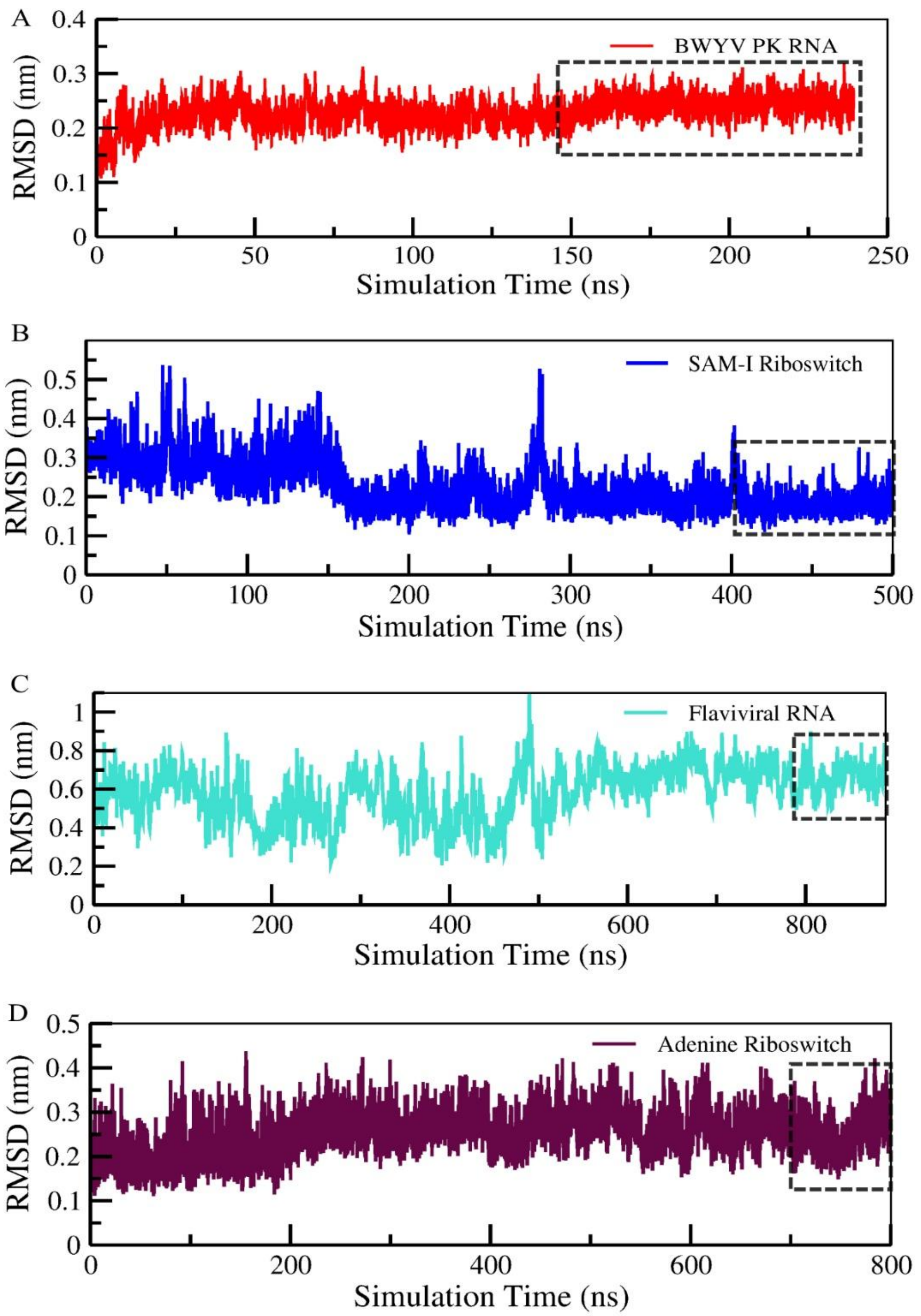


**Figure S1:** Root-mean-square deviation (RMSD) analysis of four RNA structures: BWYV PK RNA, SAM-I riboswitch, A-adenine riboswitch, and Flaviviral RNA. The highlighted regions in the figure represent the final 100 ns of the simulation, which were used for radial distribution function calculations.

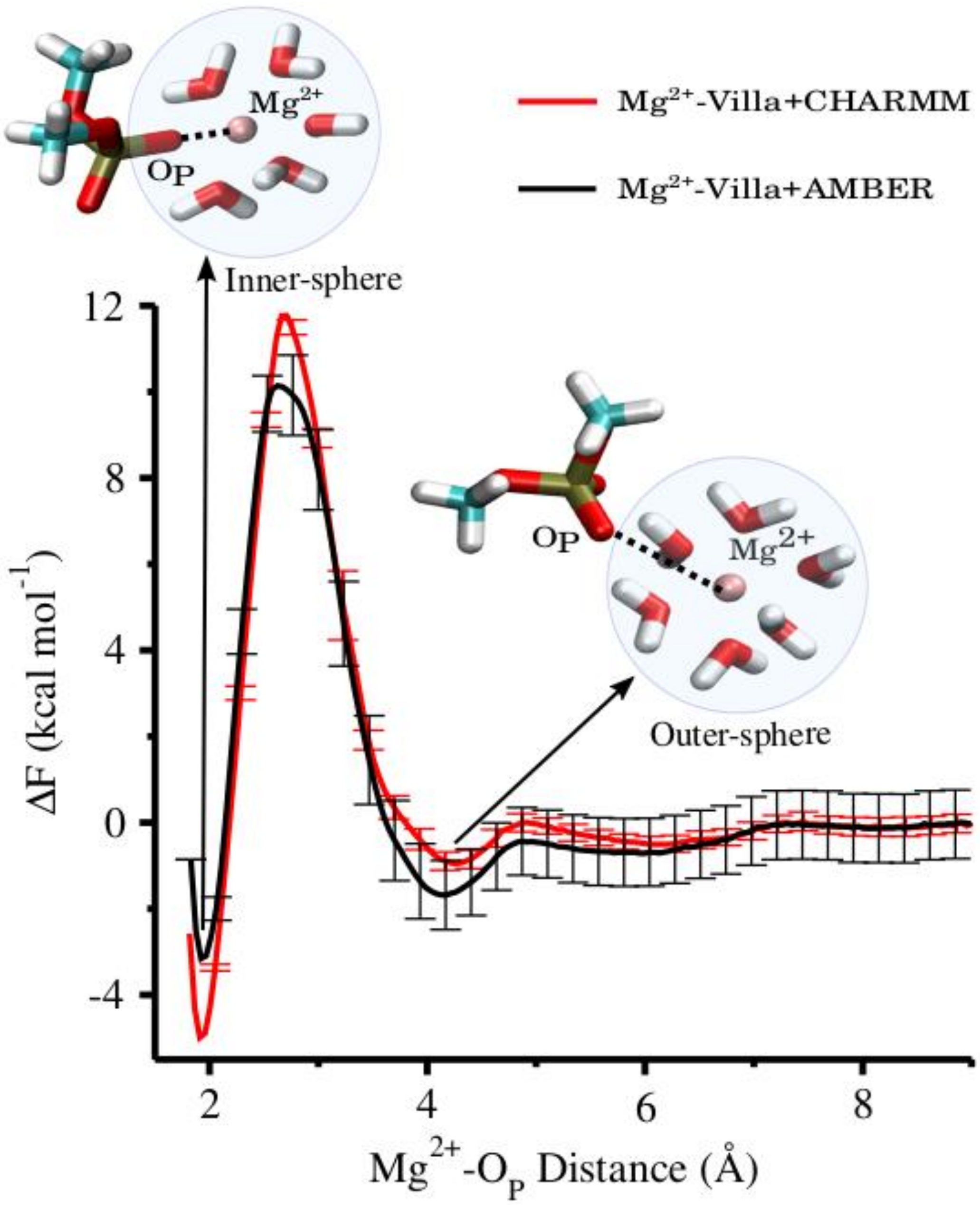


**Figure S2.** Comparison of activation free energy change during transition from inner sphere state to outer sphere state inclusion of $Mg^{2+}$-Villa parameter along a reaction coordinate ($Mg^{2+}$-$O_P$): CHARMM36 force fields (marked in red) and AMBER (amber99_parmbc0_chioL3_po) force fields (marked in black) with $Mg^{2+}$-Villa parameter were used to perform umbrella sampling. The activation barrier in CHARMM36 (16.5 kcal $mol^{-1}$) is 3.6 kcal $mol^{-1}$ is more compared to AMBER force fields (12.9 kcal $mol^{-1}$).

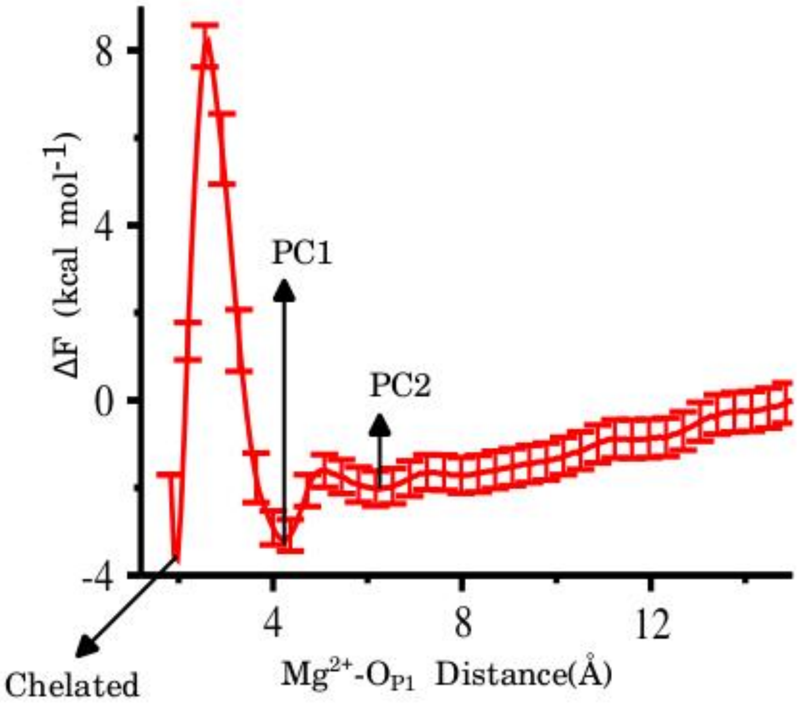


**Figure S3:** Estimation of free energy associated with unbinding of magnesium from oxygen of phosphate of a DMP molecule from chelated state to mono-coordinated prechelate1 (PC1) along a pathways ($Mg^{2+}$-$O_{P1}$ = $r_1$) and evaluated activation free energy is **12.01** kcal $mol^{-1}$. Calculated free energy was obtained by using amber99_parmbsc0_chiOL3_po force fields with incorporation of Villa $Mg^{2+}$ parameter.

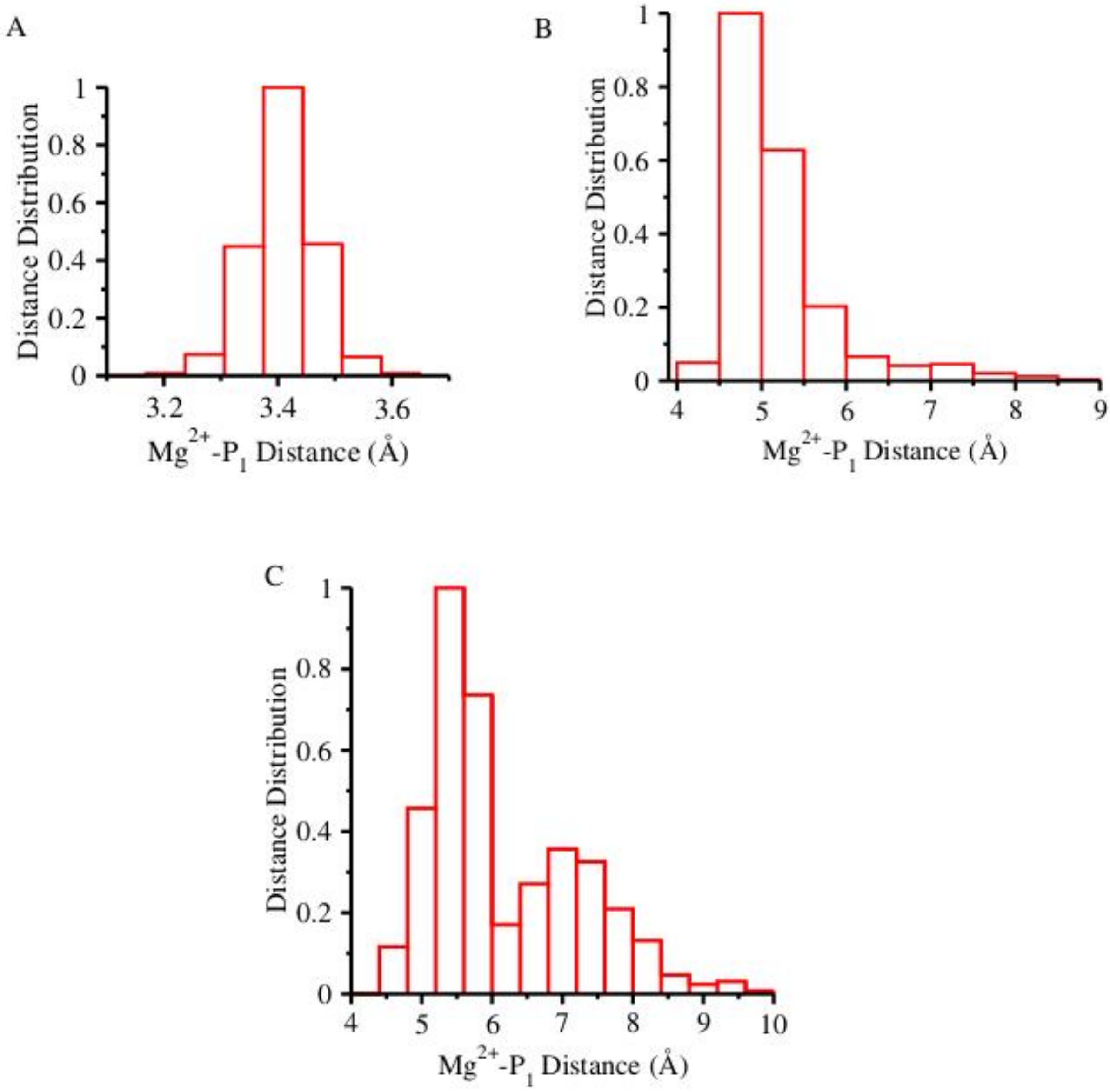


**Figure S4:** The distance distribution representation of magnesium and phosphorous of phosphate in contact layer along $r_1$ ($Mg^{2+}$-$P_1$) pathways for corresponding states of metadynamics simulations of two dimethyl phosphate molecule (DMP). (**A**) Chelated state shows distance distribution is ~3.41 Å. (**B**) The distance distribution for mono-coordinated prechelate1 (PC1) is ~4.74 Å. (**C**) While mono coordinated prechelate2 (PC2) shows two peaks ~5.39 Å and ~7.0 Å.

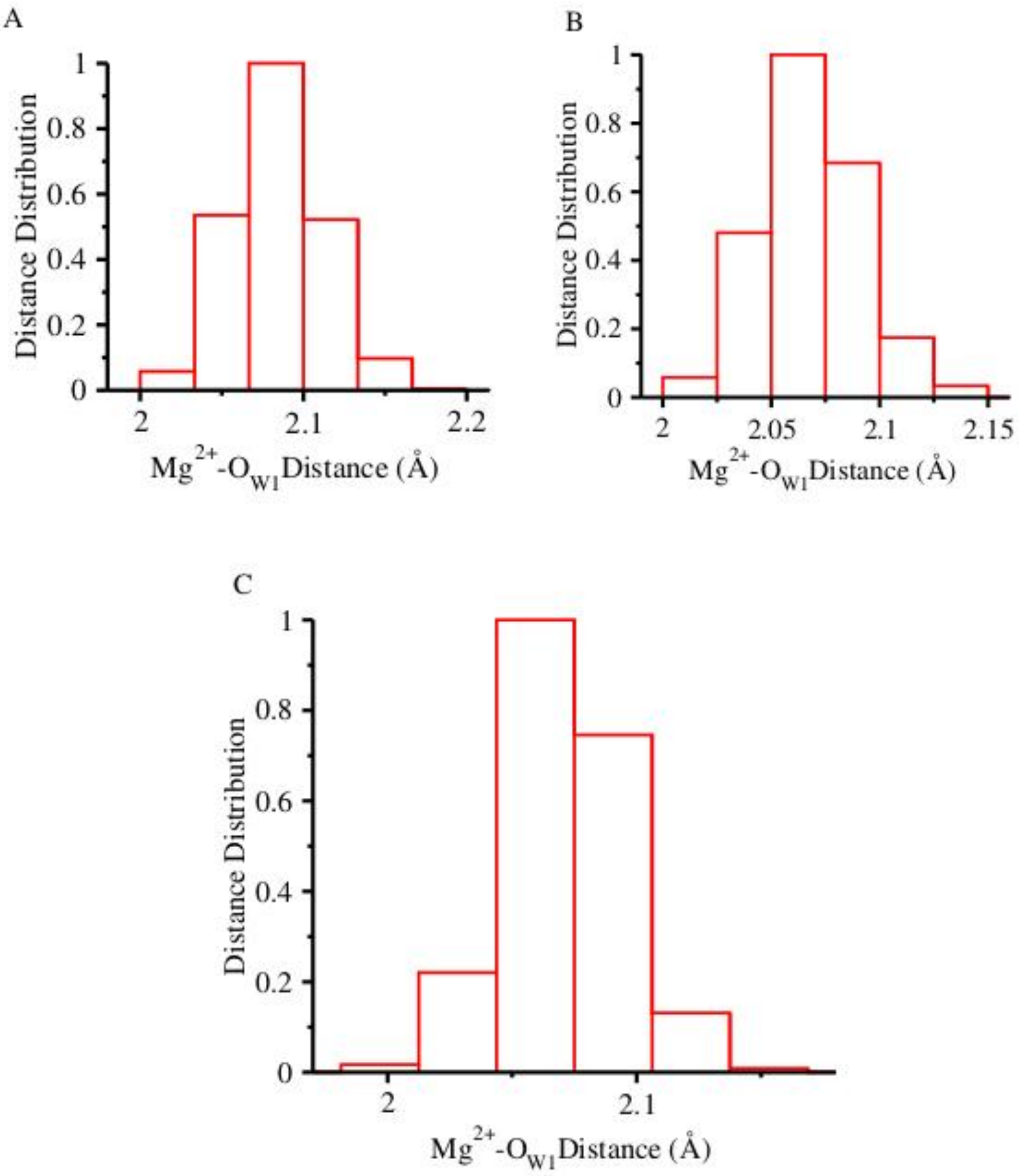


**Figure S5:** The average distance distribution representation of magnesium and oxygen of contact layer of water along $r_1$ ($Mg^{2+}$-$O_{W1}$) pathways for corresponding states of metadynamics simulations of two dimethyl phosphate molecule (DMP). (**A)** Chelated state shows distance distribution is ~2.08 Å. (**B**) The distance distribution for mono-coordinated prechelate1 (PC1) is ~ 2.07 Å. (**C**) Distance distribution of mono coordinated prechelate2 (PC2) is ~2.06 Å.

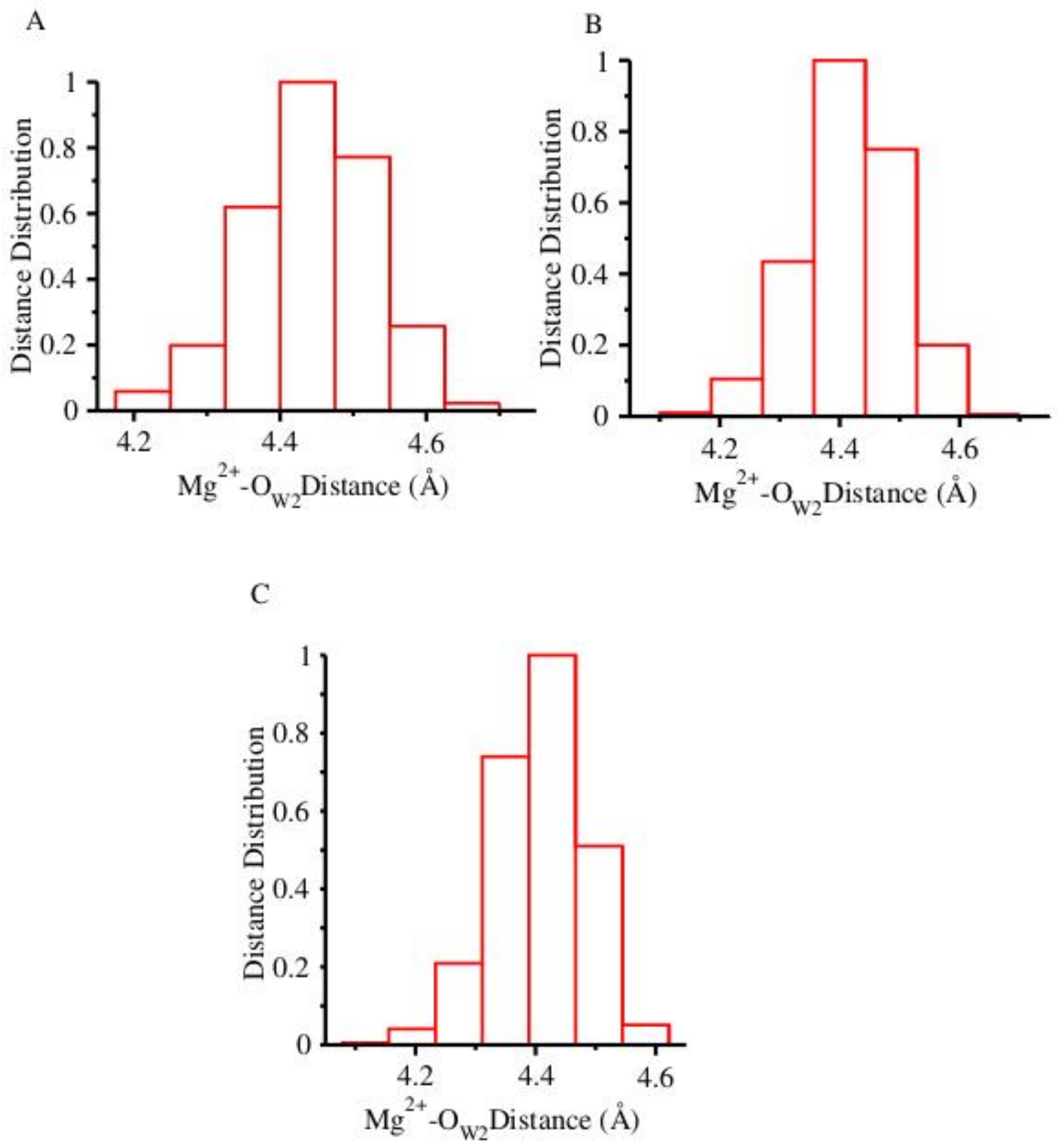


**Figure S6:** The average distance distribution representation of magnesium and oxygen of first solvent separated layer (water) along $r_1$ ($Mg^{2+}$-$O_{W2}$) pathways for corresponding states of metadynamics simulations of two dimethyl phosphate molecule (DMP). (**A)** Chelated state shows distance distribution is ~4.43 Å. (**B**) The distance distribution for mono-coordinated prechelate1 (PC1) is ~ 4.44 Å. (C) Distance distribution for mono coordinated prechelate2 (PC2) is ~4.43Å.

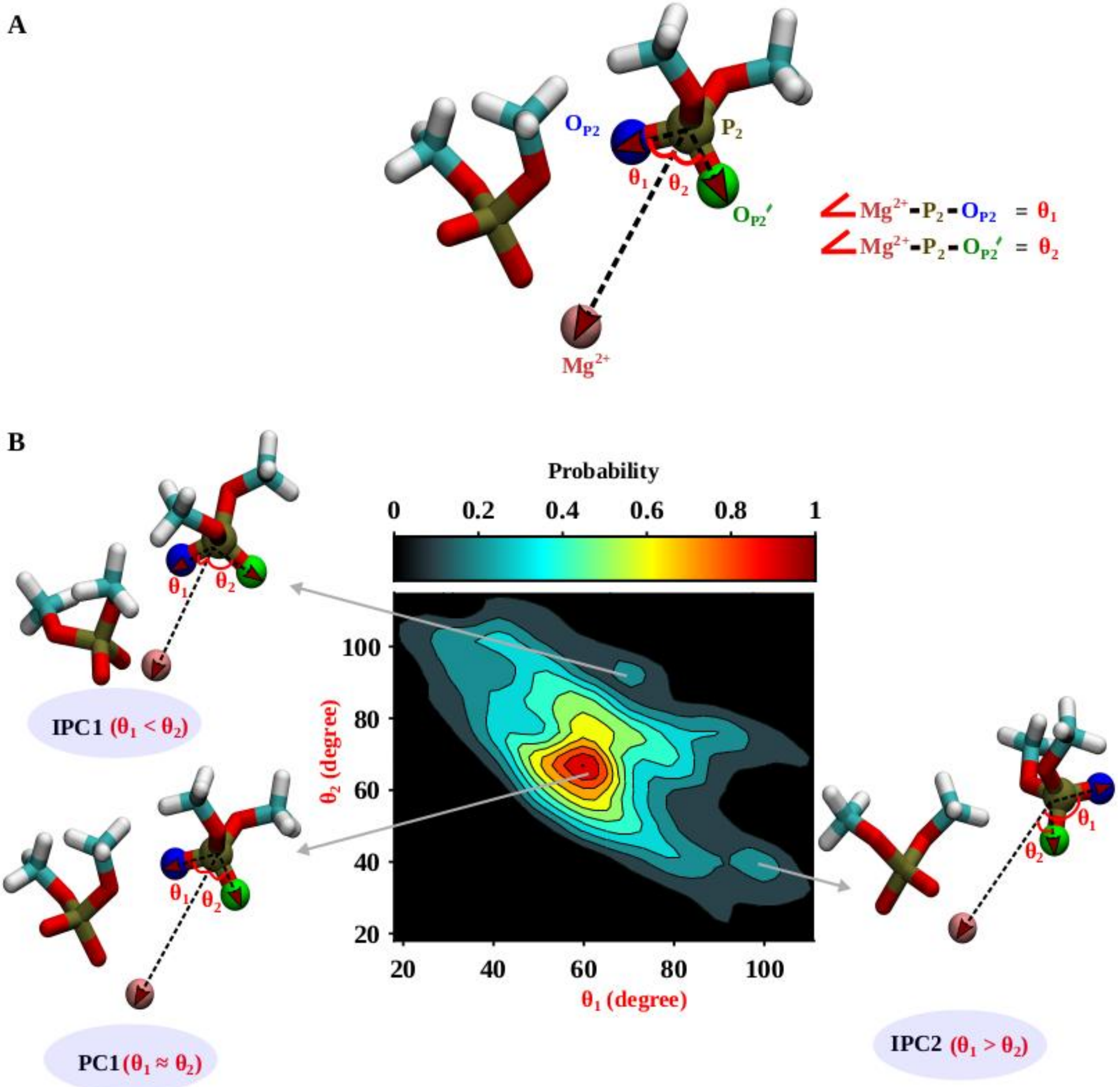


**Figure S7:** Orientational angle dependent conformations of the solvent separated phosphate relative to $Mg^{2+}$. (A) The orientational angles $\theta_1$ and $\theta_2$ are defined within the representative structure. The first phosphate oxygen $O_{P2}$ is represented in blue and the second phosphate oxygen $O_{P2'}$ is represented in green. (B) Two dimensional contour plot of probabilistic population distribution is represented as function of $\theta_1$ and $\theta_2$. The plot finds majorly three states (PC1, IPC1, IPC2) with significant probability values. The representative structures of the states are shown.

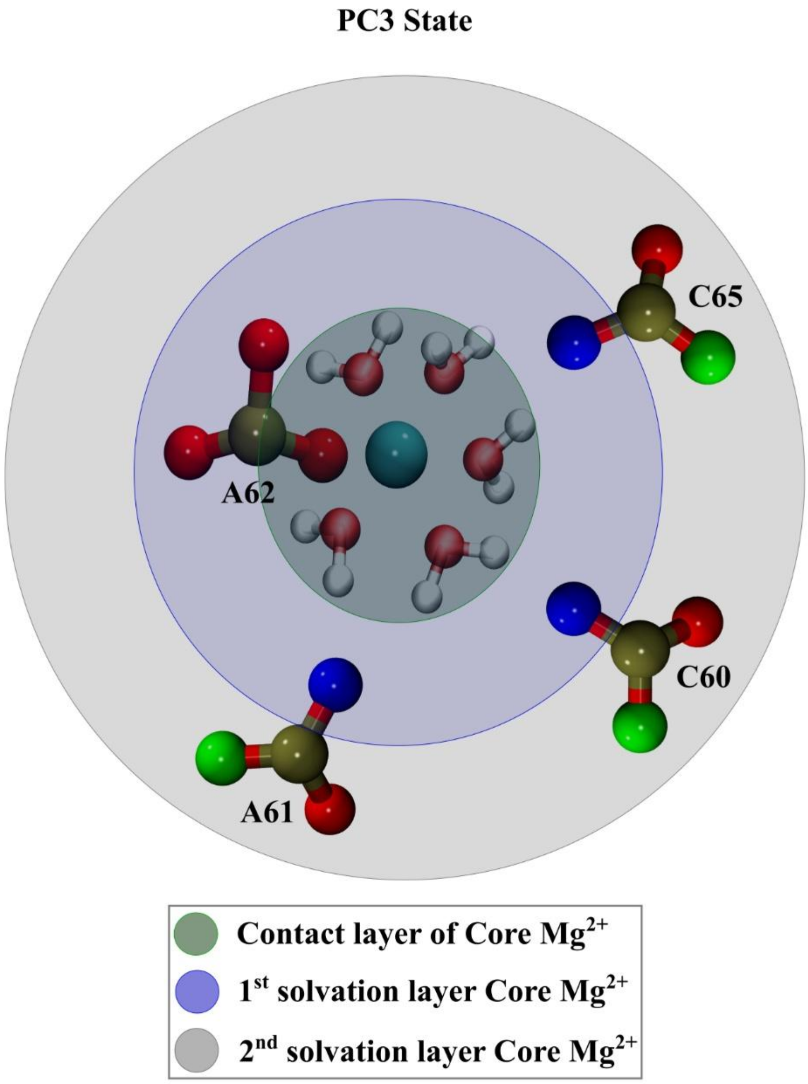


**Figure S8:** Representative structure of meta-sphere coordinated pre-chelate state 3 (PC3) of the core chelated $Mg^{2+}$ in SAM-I RNA. $Mg^{2+}$ is forming inner-sphere contact with A62 and three outer-sphere phosphates are appeared in 1st ion solvation layer.

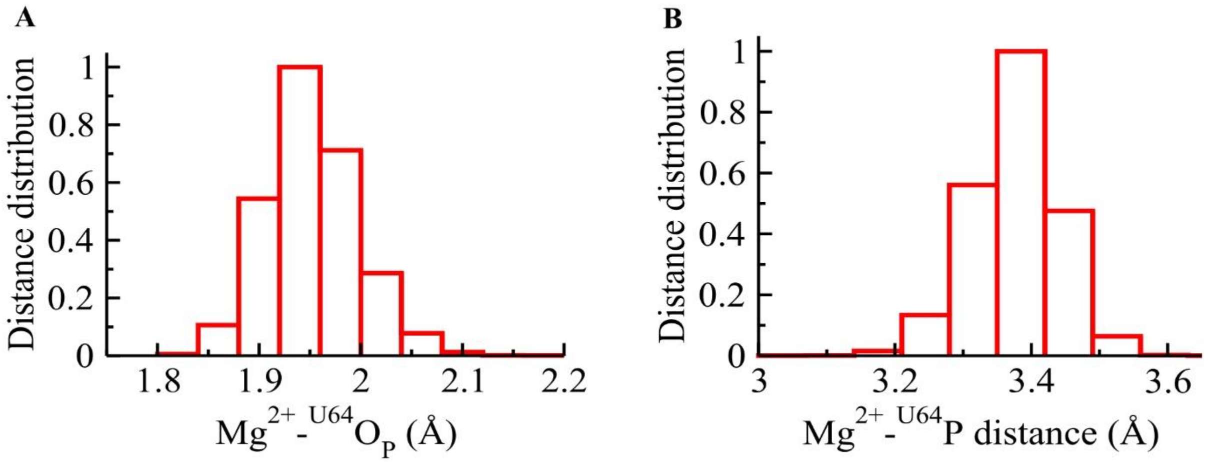


**Figure S9:** Mean distance distribution of $Mg^{2+}$-$O_P$ and $Mg^{2+}$-P in SAM-I RNA. **(A)** Distance distribution of $Mg^{2+}$ and oxygen of phosphate ($O_P$) of U64. The most probable peak appears at ~1.95 Å. **(B)** Distance distribution of $Mg^{2+}$ and phosphorus atom (P) of U64. The most probable peak appears at ~3.4 Å.
